# Unique Binding and Stabilization Mechanisms Employed By and Engineered Into Nanobodies

**DOI:** 10.1101/2023.10.22.563475

**Authors:** Natalia E. Ketaren, Peter C. Fridy, Vladimir Malashkevich, Tanmoy Sanyal, Marc Brillantes, Mary K. Thompson, Deena A. Oren, Jeffrey B. Bonanno, Andrej Šali, Steven C. Almo, Brian T. Chait, Michael P. Rout

## Abstract

Nanobodies are single domain antibody variants that bind an antigen with the precision and affinity of a conventional antibody at only a fraction of their size. In solving the crystal structures of our nanobody-GFP complexes and compared with other available structures, we uncover mechanism that enable nanobodies to function so efficiently and effectively as single-domain antibodies. We show that unlike conventional antibodies, a nanobody repertoire maximizes sampling of their antigen surface by binding a single antigen in at least three different orientations which can be predicted by their paratope composition. We also structurally reengineering these nanobodies to improve their antigen affinity, their stability, or both – results which also revealed the strong connection between nanobody stability and affinity. We achieved this by either directly modifying the paratope, or by altering a particular region within their third framework, which is a highly conserved area that we determined plays a role in controlling nanobody stability. Our study suggests that these unique characteristics of nanobodies allow them to interact with antigens as effectively as conventional antibodies, despite their smaller size. This understanding provides methods to facilitate optimizing, humanizing and functionalizing nanobodies, thus paving the way for their utilization in diverse areas such as research, diagnostics, and therapeutic development.

**Significance Statement:** Nanobodies are a unique type of antibody fragment found in select animals, containing all its antigen binding ability reduced to a single ∼15 kDa protein. There is increasing development of nanobodies for research, diagnostics, and therapeutics, yet how nanobodies function so effectively as single domain antigen binders with the precision and affinity of conventional antibodies is unclear. In this study, we present key observations to help answer this question, where one key finding is the strong relationship between nanobody stability and antigen affinity aided by the identification of a highly conserved region in nanobodies essential for maintaining nanobody stability. This region may have been retained in nanobodies in lieu of stabilizing mechanisms induced by dimerization as seen in conventional antibodies.

## Main

In nature, there exists antibody variants composed solely of IgG heavy-chain glycoproteins, rather than the heavy-chain light-chain pairs found in conventional IgG antibodies. These heavy-chain only antibodies (HCAbs), occurring in members of the family *Camelidae* (among other lineages), can bind antigens mirroring the selectivity and affinity of conventional antibodies^1^. The antigen binding domain of HCAbs is called the *V_H_H* domain, which can be independently expressed as a fragment termed a *nanobody*^1^. A nanobody, at ∼15 kDa – only one-tenth the size of conventional antibodies – can penetrate and access epitopes on antigens that are inaccessible to conventional antibodies^1,2^. Nanobodies generally have high intrinsic solubility and stability in varying environments, and their comparatively low structural complexity (including the absence of glycosylation sites) allows them to be produced in bacteria at quantity using standard laboratory techniques^3–5^. These properties make nanobodies a powerful complement technology to conventional monoclonal antibody-based research reagents, diagnostics and therapeutics^5,6^; recently gaining significant traction as promising therapeutics for the treatment of SARS-CoV-2^7–10^.

The structure of nanobodies is homologous to the variable fragment heavy chain (VH) domain of conventional antibodies, formed from a standard Ig fold with three complementarity determining regions (CDR1, CDR2 and CDR3) that typically create the paratope, separated by four framework (FR) regions that form the scaffold^11–13^. However, two notable differences distinguish nanobodies from VH domains: first, nanobodies possess on average a significantly longer CDR3 than is found in conventional antibodies^1^ and second, the FR2 region of nanobodies is more hydrophilic, correlating with the loss of dimerization with the absent variable fragment light chain (VL) domain.

Engineering and often functionalization of nanobodies is required when moving from the bench to the clinic – in particular, optimization of biophysical properties (solubility, stability etc.) and minimization of antigenicity via humanization; such humanization efforts have generally focused on mutating the more hydrophilic FR2 region to better mimic the human VH domain, although such efforts can result in loss of antigen binding or nanobody aggregation^14^. Prior studies have also explored factors that contribute to the inherent stability, specificity and affinity of nanobodies^15–19^, including, for example, the roles of intramolecular disulfide bonds^20^ and key FR residues^19,21^, however, the generality of these findings remains unclear. By solving and analyzing seven nanobody-GFP complexes and compared our findings with other available structures, we showcase the unique ways nanobodies typically maximize their accessible chemical space to bind their antigens. The structures also guided successful paratope and framework reengineering – the latter revealing the unique stabilizing role of FR3 to uncover generalizable stabilization mechanisms in nanobodies that contribute to their intrinsic stability. These findings, unique to nanobody-antigen interactions, can be readily implemented to rationally guide both nanobody humanization, and the enhancement of nanobody stability and/or affinity, particularly in the development of nanobodies as therapeutics and diagnostics.

## Results & Discussion

### Nanobodies uniquely use an expanded chemical space enabling them to maximally sample their cognate antigen surfaces

The seven nanobodies in this study originate from a single llama’s immune response to GFP, which produced a large, diverse repertoire of anti-GFP nanobodies^3^. In solving the crystal structures of these seven nanobodies in complex with GFP (Fig. 1. A, Supplementary Table 1), we observe significant sampling of the GFP surface area (Fig. 1A and B), recognizing four distinct epitope regions (groups I, II, III and IV) (Fig. 1B). Groups I, II and III corroborate the lower resolution epitope mapping of Fridy *et al*^3^, placing LaG16 and LaG43 in group I; LaG19, LaG21 and LaG41 in group II; and LaG24 in group III; in addition, a fourth epitope was mapped in this study (group IV) for LaG35 (Fig. 1B).

**Figure 1.**
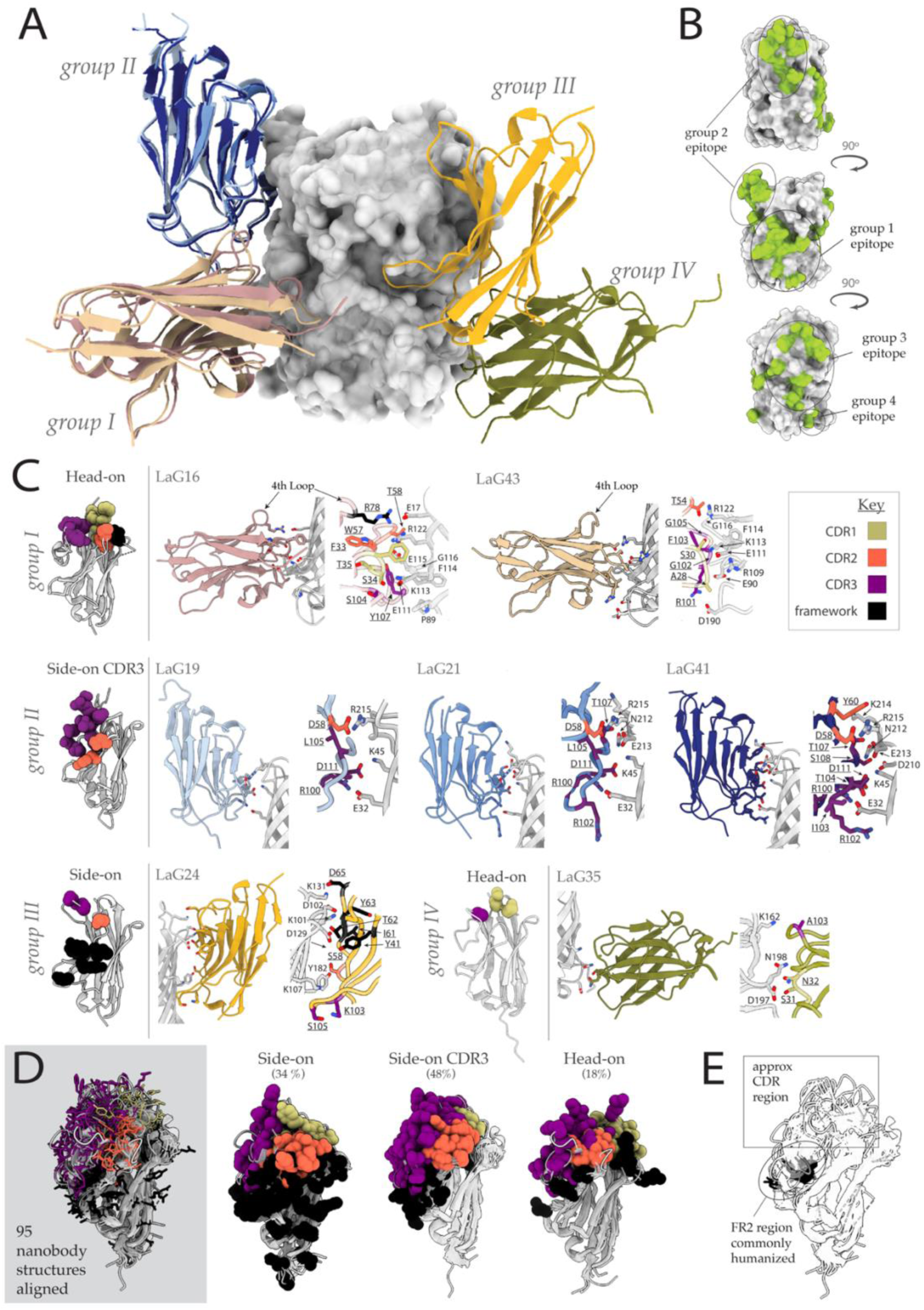
Crystal structures of 7 nanobody-GFP complexes. **(A)** The seven nanobody-GFP structures are aligned via their GFPs (in *grey*, represented as a surface rendered structure) and the nanobodies are categorized into four groups (Groups I, II, III and IV) based on their epitope on GFP. **(B)** Mapping of the four epitope groups (in *green*) on the structure of GFP (*grey*) at three different viewpoints (90° rotations). **(C)** The seven nanobody-GFP structures are categorized into their respective epitope groups, and also shown is the ‘paratope category’ they belong to: *Side-on*, *Side-on CDR3* and *Head-on,* determined by their orientation when bound to GFP. A close-up of each nanobody-GFP interaction is shown, with a detailed look at each interaction interface. The paratope residues of each nanobody are colored according to the color *key*, showing how paratope category is correlated to the contribution of different structural regions to antigen binding. **(D)** Composite of the three paratope categories highlights the degree of chemical space utilized across the 95 nanobodies analyzed, with further categorization of the 95 structures into the three paratope categories (refer to Supplementary Table 2 for PDBs). Each residue that participates in binding adopts the same color scheme as (**C**). (**E**) The same alignment of the 95 nanobodies as (**D**), highlighting amino acids (black sticks) commonly humanized within FR2, from nanobodies that use these amino acids for antigen binding.

Similar to previous studies^22–24^, the crystal structures showed how nanobodies interact in diverse orientations with GFP, dictated by the composition of their paratope (Fig. 1). This is in sharp contrast to conventional antibodies that interact with their antigens predominantly via their three CDRs in what can be referred to as a ‘Head-on’ manner (Supplementary Fig. 1). In agreement with prior work^25^ we see that the extended CDR3 loop of nanobodies, which is typically much longer than the CDR3 of the VH domain of conventional antibodies, is a dominant location for antigen binding. This is clearly observed for the three group II nanobodies, LaG19, LaG21 and LaG41 (all homologs of each other), that use their flexible CDR3 loop to mold onto the surface of GFP (Fig. 1C), biasing the orientation of the nanobodies when bound into a ‘side-on’ antigen-binding orientation. In comparison, group I nanobodies (LaG16 and LaG43) bind in a ‘Head-on’ orientation similar to conventional antibodies (Supplementary Fig. 1), where the nanobody face containing the three CDR loops is positioned to allow all three CDR loops to face and/or interact with the antigen (Fig. 1C). Both LaG43 and LaG16 interact with GFP using all three CDRs; however, only LaG16 recruits an additional “4^th^ loop” from within its framework 3 region (FR3) to interact. Remarkably, this additional loop is recruited to the paratope in numerous nanobodies^26^, where is sits next to the other CDR loops (Fig. 1C), making it in essence function as a fourth small CDR. This fourth loop has reduced size and structural diversity compared to the other three CDRs, yet contributes an essential additional binding functionality; in this sense it is analogous to the well-known additional false “panda’s thumb”^27^, again underscoring how nanobodies are able to recruit regions uncommon to classical antibody binding to form their paratopes. LaG24 (group III) also interacts in a side-on orientation with GFP, yet does so by utilizing both CDR and FR residues, with over half its paratope formed from the latter (Fig. 1C). The shorter CDR3 of LaG24 (9 residues compared to 16-23 residues for the other 6 structures, Supplementary Fig. 2) exposes its FR regions to interact (Supplementary Fig. 3), as observed with other side-on interactors (Supplementary Fig. 4). This is also illustrated by Kirchhofer *et al.*^28^, where the shorter CDR3 of their enhancer anti-GFP nanobody (also 9 residues long) enabled its FR2 to interact with GFP – the region least homologous to the human VH domain. LaG24’s FR2 and FR3 are both exposed to interact with GFP, where the latter is home to its only salt bridge with GFP (Asp65 interacts with Lys131 on GFP). This salt bridge is located at the periphery of the paratope, which may serve to secure the paratope in place. Additionally, Tyr41 within LaG24’s FR2 forms a H-bond with Asp129 on GFP. Interestingly (and concerningly), Tyr41 is one of the residues within FR2 often mutated to humanize nanobodies^14,29–31^.

Because the paratope composition of a nanobody is correlated with the orientation with which it binds its antigen, we classified our nanobodies into three broad ‘paratope categories’ (Fig. 1C): (i) Side-on interactors, where the interactions involve significant contributions from both framework and CDR regions; (ii) Side-on CDR3 – where side-on binding is due predominantly to CDR3 residues; and (iii) Head-on – where residues that form the paratope are distributed amongst the three CDRs (Fig. 1C). To investigate whether this classification system applies to other nanobody-antigen interactions, we performed a meta-analysis of 95 non-redundant nanobody-antigen complexes (Supplementary Table 1) from the Protein Data Bank (PDB, www.rcsb.org), analyzing their paratope composition and orientation when bound to their antigen using PDBePISA^32^, visual inspection via PyMoL (Schrodinger LLC 2015) and with information from their respective publications. Our analyses revealed that the three categories of interactors we deduced from our seven crystal structures could also be used to classify the 95 structures into the same three groups (Side-on, Side-on CDR3 and Head-on) (Fig. 1D). Our results revealed that ‘side-on CDR3’ interactors predominate (48%) followed by ‘side-on’ interactors (34% of the observed orientations), and ‘Head-on’ interactors (18%). The different rotations different nanobodies adopt upon antigen binding could be what enables multiple nanobodies generated against the same antigen to bind simultaneously without the effect of steric hindrance^2^. This ability endows nanobodies to participate readily in synergy^2^, where we showed previously that up to three nanobodies could bind simultaneously to a single SARS-CoV-2 spike subunit, even when their epitopes are in close proximity. Additionally, the significant participation of nanobody FR residues in antigen binding, (Fig. 1C and D), revealed several of the 95 nanobodies analyzed (∼19%) utilize their FR2 region that is often mutated for humanization purposes to bind their antigen (Fig. 1E), and of these nanobodies, all bound their antigen in the side-on or side-on CDR paratope categories. This observation underscores a potential problem with humanization efforts that target nanobody FRs, particularly FR2, as the FR can be integral for nanobody-antigen binding.

Our meta-analysis shows that our crystal structures enabled us to directly observe the structural diversity of the camelid HCAb response that maximizes antigen sampling. With FR residues playing a prominent role in binding, nanobodies targeting a single antigen bind in different orientations to their antigen, conducive to simultaneous and significant sampling of an antigen’s surface with numerous non-overlapping epitope groups (unlike conventional antibodies). This highlights how nanobodies are particularly tuned to work synergistically with other nanobodies, a phenomenon observed in our work developing therapeutic nanobodies for the treatment of the SARS-CoV-2 virus, where a pair of nanobodies can powerfully enhance each other’s interactions, to the point of their mixture being thousands of times more potent in viral neutralization than each separately^2^. Additionally, our results show that the high-frequency of FR regions that participate in antigen binding underscores potential problems when targeting FR regions to humanize or functionalize nanobodies for therapeutic and diagnostic purposes.

### Engineering to create high-affinity binders from low affinity binders

We selected two of our lower-affinity anti-GFP nanobodies to determine whether we could reengineer their paratopes to increase their affinity for GFP. Our first reengineered nanobody was a group I nanobody LaG43. Both group I nanobodies interact with their antigen in a Head-on manner with overlapping epitopes, yet only LaG16 utilizes a “4^th^ loop” on its FR3 (see above) to form its only salt bridge with GFP, between its Arg78 and Glu17 on GFP (Fig. 2A). This interaction may “lock” LaG16 into its Head-on orientation, drawing in its surface to interact with GFP, as reflected by the increased number of H-bonds at the interface (Supplementary Table. 4) and its higher affinity (0.7 nM) for GFP compared to LaG43 (11 nM). We constructed variants of LaG43 to see if we could recruit its 4^th^ loop to interact and thereby increase its affinity for GFP. We introduced an arginine at three different positions on LaG43’s 4^th^ loop, creating the following point mutants: LaG43_N78R_, LaG43_K80R_ and LaG43_N81R_. The binding affinities determined by surface plasmon resonance (SPR) revealed all LaG43 variants had increased affinity for GFP compared to wildtype (Fig. 2A, *far right*). However, while improved, the affinities of the variants for GFP did not equal that of LaG16 (Fig. 2A, *far right*). This may be due to destabilization of the salt bridge formed between Arg101 on LaG43 and Asp190 on GFP (Fig. 2A*, middle left*) by the newly introduced interaction point on LaG43’s 4^th^ loop. To determine if the interaction is between the newly introduced arginine and Glu17 of GFP, SPR experiments were performed against a GFP mutant with alanine substituting Glu17 (GFP_E17A_). The results showed restored weaker wildtype binding affinity for LaG43_N78R_ (the direct homologous mutation to LaG16’s Arg78), which supports that increased affinity is a result of an interaction between the introduced arginine and Glu17 on GFP (Supplementary Fig. 5A). However, restored wildtype binding was not observed for LaG43_K80R_ and LaG43_N81R_ against GFP_E17A_ – both showed stronger binding to GFP_E17A_ compared with wildtype GFP. It is possible that, for these point mutations, a different set of interactions formed with GFP_E17A_ resulting in increased affinity. Interestingly, thermal denaturation measurements by differential scanning fluorimetry (DSF) showed that the stability of variants LaG43_N78R_ and LaG43_N81R_ were slightly decreased (Fig. 2A, *middle right*), suggesting the charged arginine introduced at those positions may slightly destabilize the structure. This slight destabilization may allow these variants to form a better interaction interface with GFP, which may explain the correlation with increased affinity. Our results suggest recruiting the 4^th^ loop to interact in Head-on binders creates an overall better set of high-affinity LaG43 binders in the LaG43 variants over wildtype, revealing a unique mechanism to create high-affinity binders from low-affinity binders in members of the Head-on interactors.

**Figure 2.**
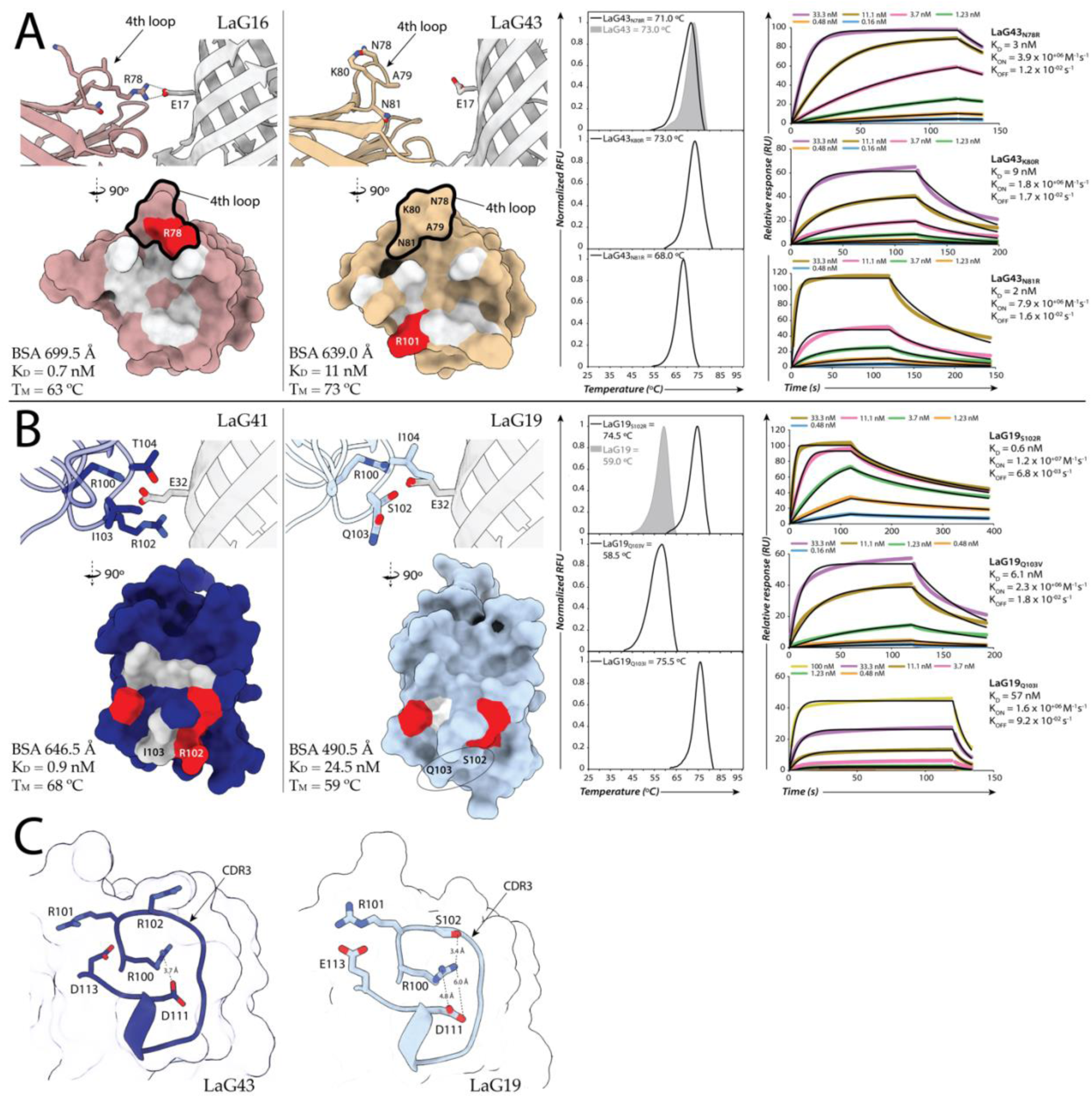
Structure guided paratope optimization. The interfaces of the nanobody-GFP complexes were analyzed using PDBePISA^40^. GFP is colored *grey*. The surface rendered structure representations are color-coded as follows: salt bridges (*red*), hydrogen bonds (*white*). **(A)** Comparison of the interaction interface of the high affinity GFP binder LaG16 (*pink*, far left) and the weaker GFP binder LaG43 (*tan*, middle left). Shown is a close-up of the 4^th^ loop region of LaG16 and LaG43, highlighting the interaction between LaG16’s Arg78 with Glu17 of GFP, the equivalent interaction is absent in LaG43. The surface rendered representation of both LaG16 and LaG43, outlines the key 4^th^ loop region (in *black*), identifying location of Arg78 on LaG16 and residues on LaG43 that were mutated to promote an interaction between its 4^th^ loop of and Glu17 on GFP. DSF results (middle right) and SPR sensograms (far right) of each LaG43 variant are plotted displaying T_M_ values and binding kinetics/K_D_ values respectively. **(B)** Comparison of the interaction interface of the strong GFP binder LaG41 (*dark blue*, far left) to the weaker GFP binder LaG19 (*light blue*, middle left). A close-up of the key LaG41 interacting residue, Arg102, and the equivalent that is absent in LaG19. A surface representation comparing the interaction interface of LaG41 and LaG19, reveals a greater hydrogen bond network in LaG41 compared to LaG19 and an extra salt-bridge that is absent in LaG19, encircled in *black*. DSF results (middle right) and SPR sensograms (far right) of each LaG19 variant are plotted displaying T_M_ values and binding kinetics/K_D_ respectively. (**C**) A detailed view of the CDR3 region surrounding amino acid 102 for LaG41 and LaG19. For LaG41, the triple arginine motif formed from Arg100, Arg101 and Arg102 repels Arg100 and Arg101 from Arg102 to promote their interaction with adjacent basic residues. For LaG19, Ser102 interacts with Arg100, with large distances between Arg100 and Asp111. The longer side-chain of Glu111 in LaG19, promotes an interaction with Arg101. The atomic distances between the interacting amino acids are represented as dashed lines with the distance in Å stated for each interaction.

The group II nanobodies have very similar structures when bound to GFP and share high sequence homology with almost identical CDR3 loops (Supplementary Fig. 2). However, LaG19 has a 27-fold weaker affinity for GFP (K_D_ = 24.9 nM) compared to the highest affinity group II nanobody LaG41 (K_D_ = 0.9 nM) (Fig. 2B, *middle left*), which we hypothesize is due to a serine at position 102 instead of an arginine as seen in the other group II nanobodies. In LaG41 and LaG21 Arg102 forms several key interactions with GFP, including an electrostatic interaction with Glu32 on GFP (Supplementary Table. 4) (Ser102 in LaG19 does not participate in binding to GFP). We created the following three point-mutants of LaG19 aimed at increasing LaG19’s affinity for GFP (Fig. 2B): LaG19_S102R_ (the key serine to arginine substitution), LaG19_Q103V_ and LaG19_Q103I_. The latter two point-mutants were designed to test whether hydrophobic amino acids promote antigen affinity, as hydrophobic amino acids are highly represented in the paratopes of nanobodies^26^. SPR of the three variants (Fig. 2B, *far right*) revealed a >40-fold increase in affinity of LaG19_S102R_ (K_D_ = 0.6 nM) for GFP compared to wildtype LaG19, making its affinity comparable to LaG41 (K_D_ =0.9 nM). We confirmed the interaction is between Arg102 in the variant and Glu32 on GFP by performing SPR with LaG19_S102R_ against a point mutant of GFP substituting Glu32 to an alanine (GFP_E32A_), revealing undetectable binding of LaG19_S102R_ (Supplementary Fig. 5B) for GFP_E32A_. Interestingly, the increased affinity for GFP was coupled to a 15 °C increase in the T_M_ of LaG19_S102R_ compared to wildtype (Fig. 2B, *middle right*). For LaG41 and LaG21, Arg102 is a member of a triple arginine motif with Arg100 and Arg101 (Fig. 2C). The concentration of like charges at this region presumably repels Arg100 and Arg101 from Arg102, bringing them closer to two adjacent acidic residues (Asp 111 and Asp113) also located on CDR3 (Fig. 2C, *left*). The repelled arginines are in a better proximity to interact with their nearby acidic residues, which may serve to stabilize the CDR3 loop, which in so doing stabilizes the overall nanobody structure resulting in the 15 °C increase in T_M_ of LaG19_S102R_. Additionally, with one of the acidic residues in LaG19 being glutamate (Glu113, Fig 2C, *right*), the longer side-chain may further promote a stable interaction with the repelled Arg101 in the LaG19_S102R_, adding to the higher stability of the variant over wildtype.

The correlation between the large increase in stability of LaG19_S102R_ with a large increase in affinity for GFP, leads us to hypothesize that the >40-fold increase in affinity is due to the cumulative effect of Arg102’s stabilizing interaction formed with Glu13 on GFP, and the intra-loop stabilization executed by the triple arginine motif of CDR3 – possibly an instance where a stabilized CDR3 promotes antigen binding. It is known that the CDR3 loops of nanobodies are important for nanobody stability and solubility, in cases where additional disulfide bonds and/or other non-covalent interactions directly stabilize these loops^15,33^, but unique here is a mechanism of intra-loop stabilization executed by the triple arginine motif and its impact on antigen affinity. Our other LaG19 point mutants allowed us to further probe the relationship between nanobody stability and affinity. LaG19_Q103I_ has a large increase in T_M_ (∼16 °C) over wildtype that is similar to LaG19_S102R_; however, unlike LaG19_S102R_, LaG19_Q103I_ has a ∼2-fold decrease in affinity for GFP (K_D_ = 57 nM) compared to wildtype. It is possible that the introduction of a hydrophobic isoleucine in this solvent-exposed region of CDR3 (Supplementary Fig. 6) creates a local region of stability on CDR3 that draws CDR3 away from the surface. This in turn stabilizes the nanobody structure (as reflected by the large increase in T_M_) yet restrains the ability of CDR3 to interact with GFP resulting in the observed decrease in affinity of the variant. Conversely, LaG19_Q103V_ has an increase in affinity for GFP over wildtype (6.1 nM), when there is no significant change in T_M_.

We demonstrate here successful reengineering of nanobody paratopes that increased antigen affinity – in both cases achieved by the optimal placement of a single arginine residue. We show successful recruitment of a “4^th^ loop” to become a part of the paratope that may prove useful when engineering the paratopes of nanobodies that also bind their antigen in a Head-on orientation. We also observe the relationship between stability and affinity in multiple. Firstly, with LaG19_S102R_, we show how increased antigen affinity is the product of two mechanisms: (i) stabilization of the epitope-paratope interface, and (ii) unique intra-stabilization of the paratope itself. Secondly, with the LaG19_Q103I_ variant, “over-stabilization” of the structure possibly restrained the nanobody paratope from binding optimally to its antigen as shown from the reduced affinity of the more stable variant. Lastly, concomitant to the previous finding, slightly *destabilizing* the nanobody paratope structure (LaG43 variants) correlated with *increased* antigen affinity, which suggests nanobodies need a degree of movement and flexibility within their paratope to optimally bind its antigen. These results reveal that the delicate balance between a nanobody’s stability and antigen affinity should be considered when directly engineering the paratopes of nanobodies.

### Key role of Framework 3 in nanobody stability

A particularly stable framework provides obvious advantages to a nanobody’s utility on the bench and the clinic. However, as shown above, we see the detrimental effect that over-stabilizing a nanobodies structure may have on its antigen binding affinity. We probed whether specific framework residues built-in to the nanobody structure can be modified to disproportionately favor framework stability without impairing affinity. When comparing the sequences of all seven nanobodies, only LaG21 lacked a highly conserved phenylalanine residue at position 69 located in FR3 at the start of its 6^th^ beta-sheet on the nanobody face opposite the CDR loops (Fig. 3A and Supplementary Fig. 2), instead having a leucine. This conserved phenylalanine forms an interaction with the sulfur group of an adjacent methionine residue (Fig. 3A, *top*). Interactions between methionine and aromatic amino acids are documented to be an important, evolutionarily conserved stabilizing interaction within protein structures^17–19^. LaG21 has high sequence and structural homology with the other group II nanobodies, but unlike LaG19, its >7-fold weaker affinity for GFP relative to LaG41 cannot be increased via direct paratope optimization, as its CDR3 loop is almost identical to LaG41 (Supplementary Fig. 2). Additionally, LaG21 has slower association with GFP (K_on_) and a markedly fast dissociation rate (K_off_)^3^, which coupled to it having the lowest T_M_ of all seven nanobodies at 55°C (Fig. 3A), suggests that the greater motion in the structure of LaG21 may affect its antigen binding. It is possible that the absence of this critical Phe-Met interaction affects the overall stability of the nanobody that in turn affects binding to GFP, so we created a LaG21 variant by substituting Leu69 with phenylalanine (LaG21_L69F_) to hopefully increase stability and potentially also, affinity. Our DSF results revealed the LaG21_L69F_ variant indeed increased in structural stability, with a 9°C increase in T_M_ compared to wildtype (Fig. 3A, *middle*). Additionally, SPR revealed a 10-fold increase in affinity of LaG21_L69F_ (0.7 nM) for GFP over wildtype LaG21 (7 nM), bringing its affinity on par with the strongest binder to GFP of the group II nanobodies (LaG41). This suggests that the highly conserved Phe69 is important for antigen binding via framework stabilization, due to its stabilizing interaction formed with its adjacent methionine (Met84 in group II nanobodies).

**Figure 3.**
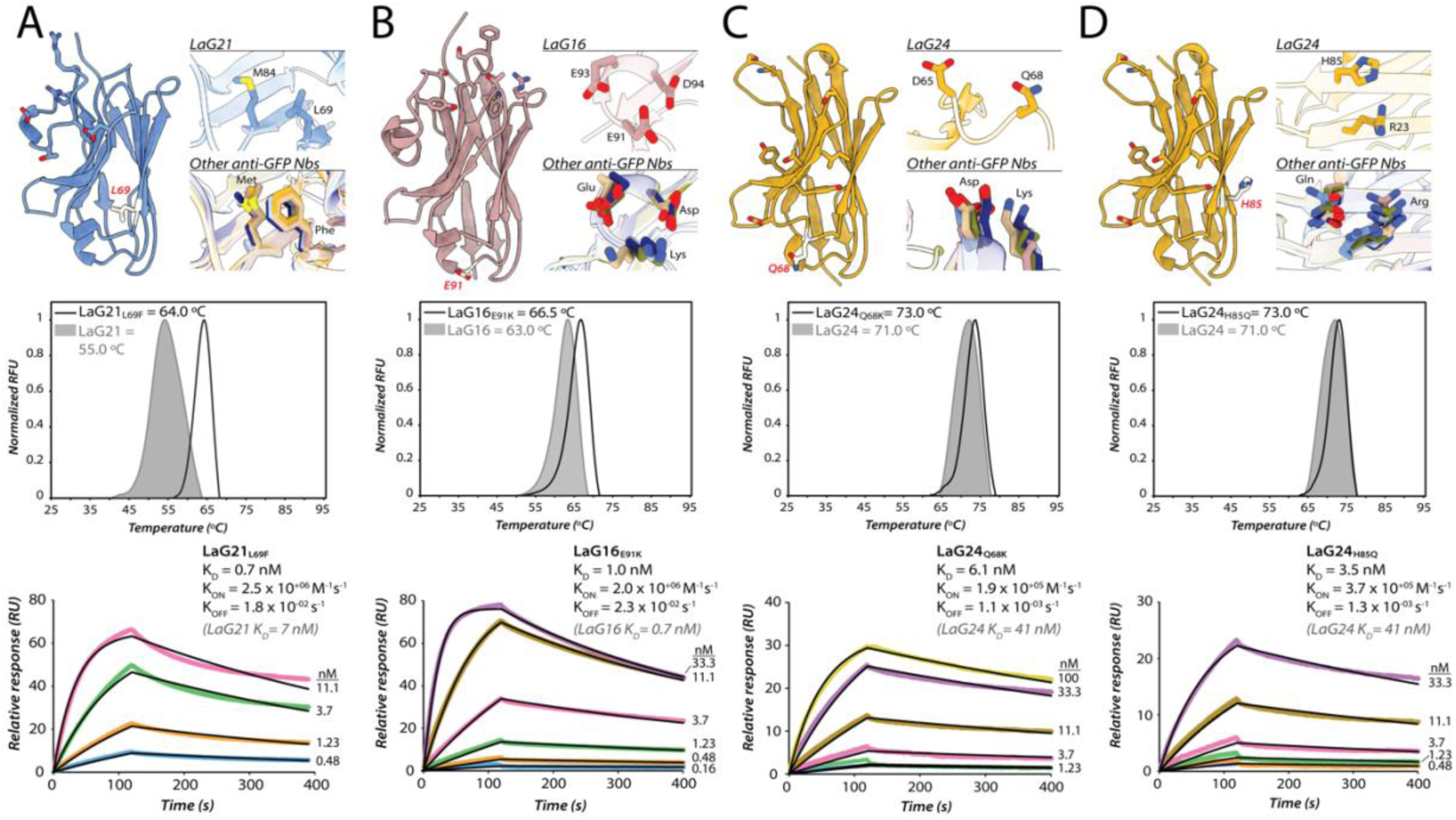
Framework optimization of anti-GFP nanobodies. **A-D** shows the rationale behind each FW optimizing mutation and the results of their characterization using DSF and SPR revealing T_M_ and kinetic/K_D_ values compared to wildtype. The position of each mutated FW residue is colored *white* and represented in *stick form* on the structure of the respective nanobody. The paratope of each nanobody is also shown, represented as sticks. (**A**) LaG21_L69F_ rationale, with close-ups of LaG21’s highly conserved methionine (Met84) pointed away from Leu69 and the equivalent residues in the other six anti-GFP nanobodies where Phe is in close proximity to the sulfur group of the highly conserved methionine. (**B**) LaG16_E91K_ rationale, showing a detailed view of Glu91 of LaG16 flanked by two conserved acidic residues (Asp94 and Glu91) in comparison to Lys91 in the other six anti-GFP nanobodies flanked by the same highly conserved acidic residues. (**C**) LaG24_Q68K_ rationale, with close-ups of Gln68 of LaG24 flipped away from the acidic Asp65 unlike in the other six nanobodies, where the basic lysine residue is oriented towards the acidic aspartate residue. (**D**) LaG24_H85Q_ rationale, with close-ups of Arg23 on LaG24 flipped away from His85, in comparison to the interaction of Gln85 with arginine present in the other six anti-GFP nanobodies.

We identified three additional sites on our anti-GFP nanobodies that deviate from the consensus FR sequence with stabilizing roles (Fig. 3B – D). The first site, Glu91 on LaG16, is located on a loop also within FR3 that again is positioned on the opposite face of the CDR loops; Glu91 is flanked on either side by two acidic amino acids (Asp94 and Glu93) (Fig. 3B, *top*). The clustering of like charges increases the likelihood of coulombic repulsion, known to decrease protein stability^20^. In the other six nanobodies, a lysine is present in place of this glutamate, also flanked by two acidic amino acids. In contrast to Glu91, the lysine would neutralize the effect of charge repulsion, creating local stabilization within the loop. To test this possible stabilizing effect, we constructed a LaG16_E91K_ variant and assessed its stability and binding (Fig. 3B). Our DSF data showed the T_M_ of the variant did indeed increase by 3°C compared to wildtype, without impairing affinity (Fig. 3B and Supplementary Table 3). The second location identified was Gln68 of LaG24 (this residue has also shown to be stabilizing in a previous study^19^), located on a loop within FR3 that is in a similar region to the other two identified sites (Fig. 3C). This residue is a lysine in the other six nanobodies, and this lysine interacts with a nearby conserved aspartate residue (Fig. 3C). We hypothesize that the interaction between lysine and aspartate serves to stabilize this loop region. We constructed and tested a LaG24 point mutant substituting glutamine with lysine, LaG24_Q68K_. Our DSF results revealed a 2°C increase in thermal stability coupled to an almost 10-fold increase in affinity of LaG24_Q68K_ over wildtype (Fig. 3C and Supplementary Table 3). The final location identified was His85 on LaG24, which in the other six nanobodies is a glutamine that forms a key interaction with a conserved Arg23 located within FR1 (Fig. 3D). The location of His85 on the 7^th^ beta-sheet of LaG24 is within a similar region on FR3 as the other sites identified, and near a loop in FR3. In LaG24, the side-chains of His85 and Arg23 are flipped away from each other (Fig. 3D), where His85 interacts instead with Ser74. We created a LaG24 variant (LaG24_H85Q_), substituting His85 with glutamine to create an interaction partner for Arg23 analogous to the other six nanobodies. Similar to the other three sites, LaG24_H85Q_ had an increase in thermal stability (by 2°C) coupled to a >10-fold increase in affinity for GFP (Fig 3D and Supplementary Table 3).

Here we have identified four sites within a localized region of the FR3 of nanobodies distal to their CDR loops and not directly involved in antigen binding, which is critical for nanobody stability, and that can also be tuned to increase binding affinity (Fig. 3 and Supplementary Fig. 7). Interestingly, these four stabilizing residues are found on or next to loops, which likely constrains the dynamics of these loops to in turn stabilize the overall nanobody structure – suggesting the flexibility of loops on nanobodies influences their stability and consequently their antigen affinity. These four sites we identified are quite conserved in nanobodies generated against different antigens and isolated from different camelid species (though with significant remaining variation) (Fig. 4A) yet are less conserved in the VH domain of conventional antibodies (Fig. 4B). Nanobodies may have retained these localized stabilizing interactions to serve as a critical stabilization hub that reinforces the nanobody scaffold, which may compensate for the loss of VH-VL dimerization that induces a stabilizing effect in conventional IgGs. As such, these four sites embedded in the FR3 of nanobodies can be readily screened for, via primary sequence alone, to rapidly optimize nanobody stability and potentially also be used to tune nanobody affinity.

**Figure 4.**
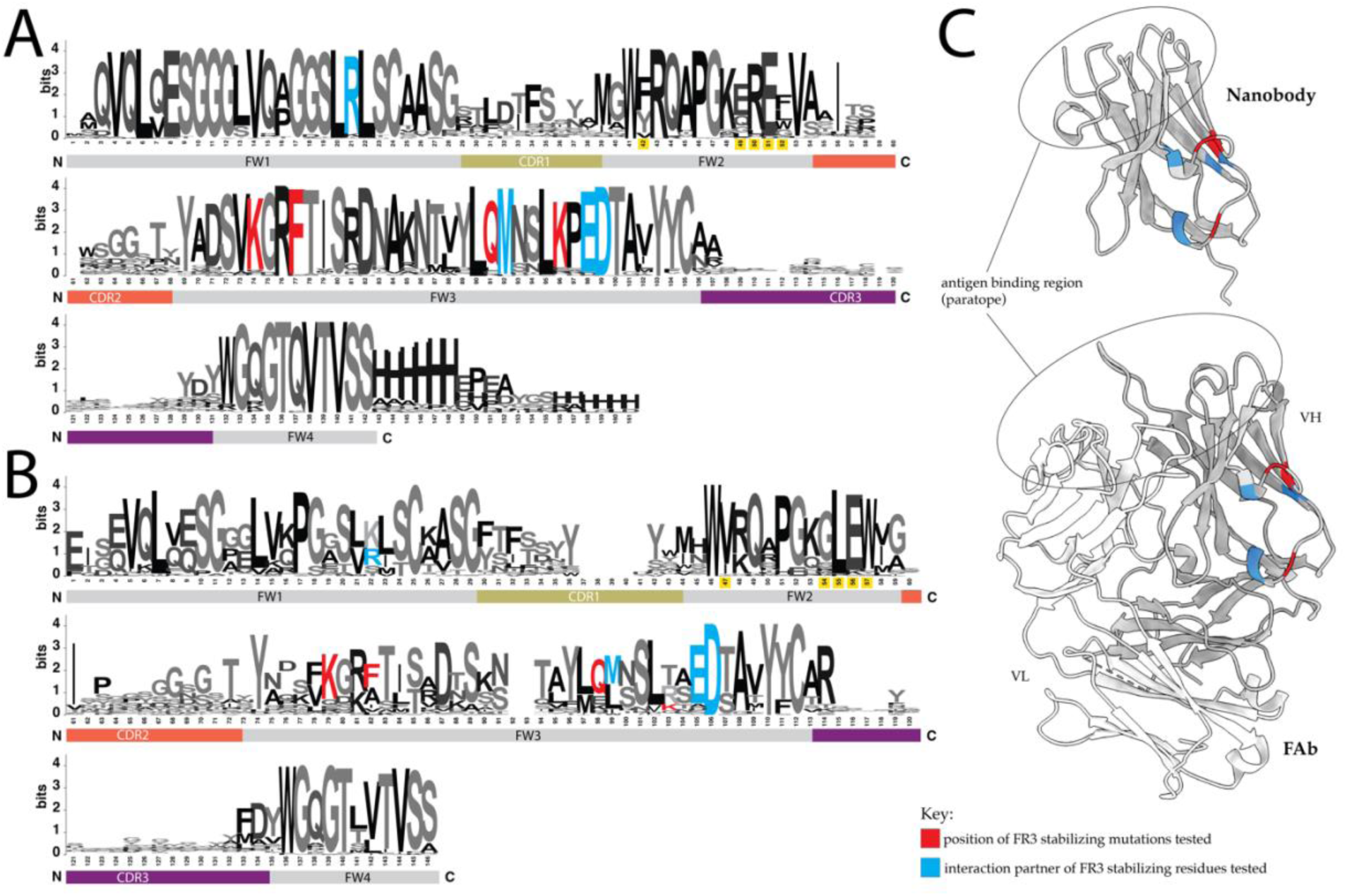
Sequence alignment of 95 nanobody sequences (refer to Supplementary Table 2 for PDBs) (**A**) and 87 VH domain sequences (refer to Supplementary Table 3 for PDBs) (**B**). The FW residues mutated and identified as stabilizing for nanobodies are colored according to the displayed key. The FW2 residues that differ between nanobodies/VHH and VH, and are commonly humanized have their sequence number shaded in yellow on each alignment. (**C**) The structure of a representative nanobody (top, LaG41 structure) and FAb (PDB ID: 1bj1), highlighting the location of the nanobody FR3 stabilizing amino acids, which are colored according to the displayed key.

### Applying general framework 3 optimization principles to nanobodies with therapeutic potential

We tested the generalizability of our identified conserved mechanisms of framework stabilization identified above on a repertoire of anti-SARS-CoV-2 spike S1 nanobodies. These nanobodies were generated from our previous work^2^ and were isolated from a different llama from which the anti-GFP nanobodies were generated. We had no atomic-scale structural information for these nanobodies, so we thus also tested the ability of these stabilizing principles to be applied with only primary sequence-level information.

In total, we constructed eight variants of eight different SARS-CoV-2 anti-spike S1 nanobodies (Supplementary Table 8), and the results of their characterization are summarized in Table 1. Importantly, the general trend is indeed that these mutations result in both increased *stability* and *affinity*. For example, the nanobody S1-40, with an affinity to spike protein too low to obtain a reliable SPR result (likely sub-micromolar K_D_), was a candidate for the introduction of a lysine at residue 89 to reduce coulombic repulsion (based on LaG16_E91K_ seen above). The resulting variant S1-40_S89K_ displayed a ∼2°C increase in T_M_ over wildtype, coupled to a remarkable 0.5 nM affinity for spike. This substitution not only “rescued” the binding ability of the nanobody but also created a high affinity binder from a nanobody with formerly no detectable binding. Similarly, creating the same stabilization site on nanobody S1-RBD-42 (to create the point mutant S1-RBD-42_E88K_) resulted in both increased stabilization and affinity for spike over wildtype (Table 1). One exception to the trend of increased affinity with increased structure stability is when the Phe-Met stabilizing interaction was introduced into an already high-affinity binder, S1-36 (K_D_ = 0.2 nM). The S1-36_L70F_ variant, similar to LaG21_L69F_, resulted in a large (∼10°C) increase in T_M_ over wildtype; however, unlike LaG21_L69F_, S1-36_L70F_ loses its ability to bind spike S1. The dramatic stabilizing effect (likely via the Phe-Met interaction) observed for S1-36_L70F_, is another example of “over-stabilizing” the structure (see above), which underscores that some empirical experimentation with several variants is important for performance improvement of a given nanobody. But also suggests that there is a stability threshold that has been hit on these over-stabilized nanobodies that disrupts the equilibrium between binding and stability. The only nanobody variant to exhibit a decrease in thermal stability was S1-RBD-14_E89K_, which had a ∼3°C decrease in T_M_ compared with wildtype. The substitution introduced a lysine in FR3 in place of a glutamate (E89K), which was predicted to stabilize this loop region within FR3 by reducing coulombic repulsion. However, with this decrease in T_M_ came a dramatic ∼80-fold increase in affinity of the S1-RBD-14_E89K_ (K_D_ = 0.3 nM) over wildtype (K_D_ = 25 nM). This is a pronounced example of the earlier observation of how slightly destabilizing the nanobody structure can enable a nanobody to bind more efficiently to its antigen, in this case to spike S1.

**Table 1.**
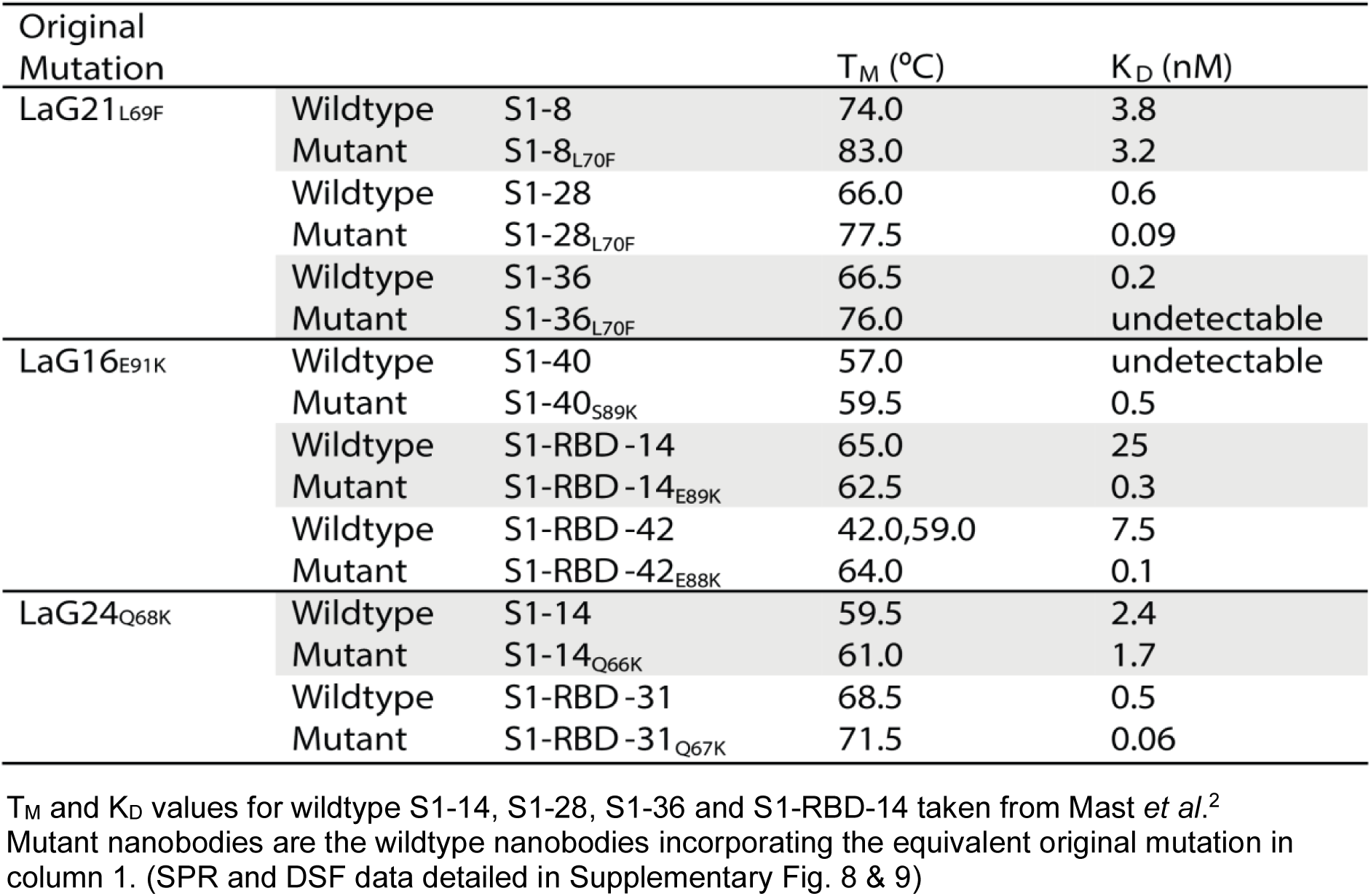
Framework optimization of nanobodies generated against wildtype SARS-CoV-2 spike protein.

| Original Mutation |  |  | T<sub>M</sub> (°C) | K<sub>D</sub> (nM) |
| --- | --- | --- | --- | --- |
| LaG21<sub>L69F</sub> | Wildtype | S1-8 | 74.0 | 3.8 |
|  | Mutant | S1-8<sub>L70F</sub> | 83.0 | 3.2 |
|  | Wildtype | S1-28 | 66.0 | 0.6 |
|  | Mutant | S1-28<sub>L70F</sub> | 77.5 | 0.09 |
|  | Wildtype | S1-36 | 66.5 | 0.2 |
|  | Mutant | S1-36<sub>L70F</sub> | 76.0 | undetectable |
| LaG16<sub>E91K</sub> | Wildtype | S1-40 | 57.0 | undetectable |
|  | Mutant | S1-40<sub>S89K</sub> | 59.5 | 0.5 |
|  | Wildtype | S1-RBD-14 | 65.0 | 25 |
|  | Mutant | S1-RBD-14<sub>E89K</sub> | 62.5 | 0.3 |
|  | Wildtype | S1-RBD-42 | 42.0,59.0 | 7.5 |
|  | Mutant | S1-RBD-42<sub>E88K</sub> | 64.0 | 0.1 |
| LaG24<sub>Q68K</sub> | Wildtype | S1-14 | 59.5 | 2.4 |
|  | Mutant | S1-14<sub>Q66K</sub> | 61.0 | 1.7 |
|  | Wildtype | S1-RBD-31 | 68.5 | 0.5 |
|  | Mutant | S1-RBD-31<sub>Q67K</sub> | 71.5 | 0.06 |

T<sub>M</sub> and K<sub>D</sub> values for wildtype S1-14, S1-28, S1-36 and S1-RBD-14 taken from Mast *et al.*<sup>2</sup>

Our results here show that the stabilization sites we identified on our anti-GFP nanobodies exert similar stabilization effects on our anti-SARS-CoV-2 spike S1 nanobodies, underscoring the generalizable nature of utilizing this region to manipulate nanobody stability and affinity. We underscore earlier observations that nanobody stability and affinity are linked, showing the tunability of nanobody affinity by targeting these stabilizing interactions. Importantly, our results further support our earlier observation that the FR3 region is a stabilization core on nanobodies – home to a number of key stabilizing interactions revealing the possible role of this FR3 region in imbuing nanobodies with the stability needed to bind an antigen with the precision and affinity of conventional antibodies in lieu of VH-VL dimerization. As such, these findings should be considered in future functionalization of nanobodies.

### Molecular modeling provides further mechanistic insight into the effect of framework 3 stabilization on nanobody function

To assess the structural differences resulting from the Leu69Phe substitution in LaG21, we performed molecular dynamics (MD) simulations of wildtype LaG21 and the FR optimized LaG21 (LaG21_L69F_). The MD simulations were performed at three temperatures beginning at ∼room temperature (300 K) then 336 K and 421 K, where the last serves as a control reflecting expected thermalization and unfolding for both the wildtype and FR optimized LaG21. Our original hypothesis (see above) is that the Leu69Phe substitution results in an interaction of the Phe with an adjacent methionine that stabilizes FR3, which in turn stabilizes the overall structure of the nanobody resulting in better binding to GFP (Fig. 3A). The MD simulations revealed that LaG21 has an overall greater degree of motion in its structure compared to LaG21_L69F_ (Fig. 5A). This degree of fluctuation is greater for residues within the CDRs, especially CDR3, which is the main interaction point of LaG21 with GFP (Fig. 1C). In comparison, LaG21_L69F_ consistently maintains low RMS fluctuations across its whole structure (Fig. 5). To probe the cause of the differences in the structural dynamics between LaG21 and LaG21_L69F_, we performed Mutual Information (MI) analysis^34^ on the MD simulations, which allows us to observe correlated dynamics between backbone dihedral angles of residue pairs in the wildtype and mutant LaG21 structures. A higher MI between residues indicates that their backbone conformations are correlated while a lower value indicates that their motions are nearly independent of each other. Our results reveal that the wildtype LaG21 shows a high degree of correlation between residue pairs in general, with significant correlations between CDR3 and FR2 and between CDR3 and the beginning of FR3, which is the location of the Leu69Phe substitution (Supplementary Fig. 10). Therefore, it is likely that the FR3 dynamics of LaG21 directly drive structural changes in its paratope, while the Leu69Phe mutation suppresses such an influence. These results support our hypothesis that stabilizing FR3 is indeed stabilizing the overall structure of LaG21, due to FR3’s direct influence on CDR3, where the decreased fluctuations of LaG21_L69F_’s CDR3 likely promotes favorable binding to GFP, resulting in the observed high affinity interaction with GFP. This hypothesis is consistent with the importance of the stabilization of CDR3 for the interaction of other group II, side-on CDR3 binders (see above example with LaG41 and LaG19).

**Figure 5:**
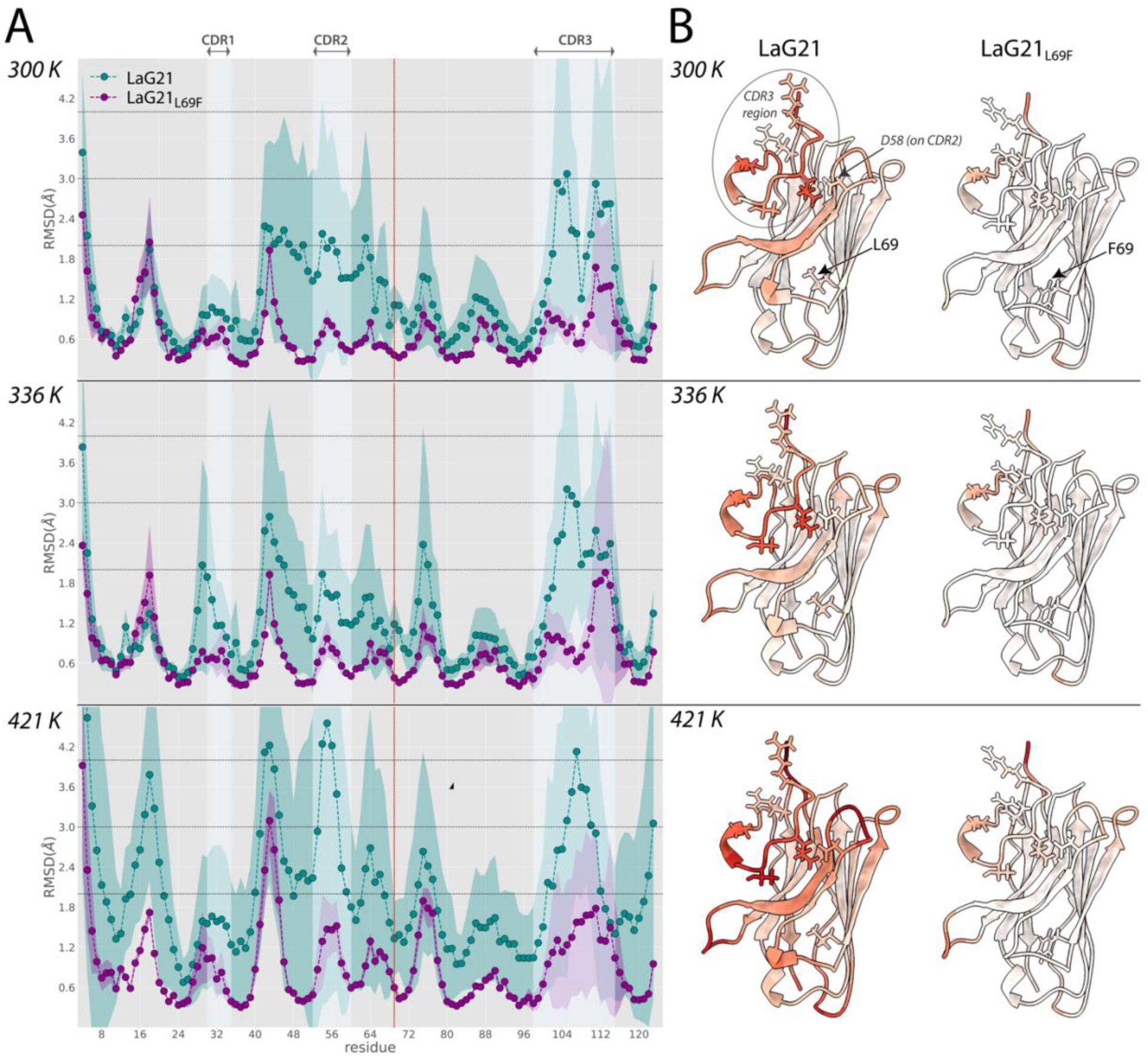
Molecular modeling of the conformal heterogeneity of LaG21. **A** and **B** present structural metrics computed from 1.6 µs molecular dynamics simulations of solvated LaG21 (in *teal*) and LaG21_L69F_ (in *purple*) at three different temperatures 300, 336 and 421 K where the shaded regions show 95% confidence intervals. **A**) The red vertical line denotes the location of the Leu69Phe mutation. In addition to CDR regions being labelled, FR and CDR regions are shaded vertically in *grey* and *light blue* respectively. Average RMS fluctuations per residue, relative to the energy minimized crystal structure at room temperature: both LaG21 and LaG21_L69F_ consistently maintain 1.8-2 Å variation in CDRs 1 and 2 at low to moderate temperatures. LaG21 has a higher baseline RMSD in CDR3 and completely unfolds at higher simulation temperatures unlike LaG21_L69F_. (**B**) Paratope residues are represented as sticks in addition to Leu69 (for LaG21 structures) and Phe69 (LaG21_L69F_ structures). Per-residue RMS fluctuations at different temperatures normalized as a Z-score between 0-100 and projected on reference LaG21 and LaG21_L69F_ structures. At near room temperature (300 K), LaG21 paratope residues Arg102, Leu105, Thr107, Asp111 show the most significant fluctuation from the crystal structure, while the Leu69Phe mutation makes the entire LaG21_L69F_ CDR3 more stable, in addition to FR regions.

In conclusion, our analysis underscores the importance of stabilizing the CDR3 when it forms the majority of the paratope, where reducing the dynamics of CDR3 likely promotes a stable interaction between CDR3 and its antigen. Molecular modeling reveals a direct correlation between FR3 and overall nanobody stabilization, which, in this case directly impacts nanobody affinity. These results underscore our finding that FR3 serves as a key stabilization core for nanobodies.

### Summary and Perspectives

Our results shed light on how nanobodies exploit their small structure and biochemical makeup to function as highly effective single-domain antigen binders, utilizing both their increased surface hydrophilicity and intrinsic stability. The ability of nanobodies to bind an antigen in different orientations, coupled with their small size, maximizes antigen surface sampling and likely enables multiple nanobodies to bind a single antigen simultaneously^2^, increasing the potential for synergy between different HCAbs during an immune response – a phenomenon showcasing the uniqueness of the HCAb immune response in animals such as camelids with this dichotomous immune system. Our paratope and framework reengineering results underscore how the structural stability of nanobodies influences a nanobody’s affinity for its antigen. Furthermore, our FR reengineering enabled us to pinpoint a region on nanobodies within FR3, distal to the CDR loops, that acts as a “stabilization core”, formed from several key stabilizing interactions peppered throughout the region. Our MD simulations rationalize this observation by suggesting that atomistic fluctuations in the FR3 region have a direct influence on CDR3 loop dynamics, which can then be modulated through FR3 re-engineering such as the point mutations we show. The stabilizing interactions found within FR3 are strongly conserved in all nanobodies in contrast to antibody VH domains, which suggests nanobodies have evolved to retain these integral stabilizing interactions – potentially as a substitute for the stabilizing effect of adjacent protein subunits found in conventional IgG antibodies. This nanobody feature allows guided optimization of nanobodies using only their primary sequence, as demonstrated by our affinity and stability enhancement of potentially therapeutic nanobodies against SARS-CoV-2 with a single point mutation. With the growing interest in utilizing nanobody technology^35–41^, we hope this study will aid researchers to maximally utilize their nanobody repertoires and rationally engineer them, including for use as diagnostics, prophylactics and therapeutics.

## Acknowledgements

We would like to thank the Fisher Drug Discovery Resource Center (DDRC) (RRID:SCR_020985) at Rockefeller University, and especially former members Lavoisier Ramos-Espiritu and Carolina Adura for technical and data analysis support as well as the Rockefeller Structural Biology Resource Center (RRID: SCR_017732). We also thank members of the Rout, Almo and Chait labs for support and assistance. We acknowledge the support of the G. Harold and Leila Y. Mathers Charitable Foundation, the Robertson Therapeutic Development Fund, the Bay Area Lyme Foundation, the Jain Foundation, and the National Institutes of Health grants P41 GM109824 (M.P.R., B.T.C. and A.S.), R21 AI154180, U54 GM094662 (S.C.A.), U01 GM098256 (M.P.R. and A.S.), and R01 GM083960 (A.S.). Data was collected at NSLS beamline X29A that was funded by the Offices of Biological and Environmental Research and of Basic Energy Sciences of the US Department of Energy (P41 RR012408), and from the National Center for Research Resources of the National Institutes of Health (P41 GM103473).

## Methods

### Anti-GFP Nanobody Generation

Anti-GFP nanobodies were generated as previously described^3^. In brief, we immunized a llama with GFP. After a strong immune response was elicited, we collected lymphocytes from bone marrow, highly enriched for active B cells. We then purified RNA from these cells and performed high-throughput sequencing on the PCR-amplified variable region (VHH) of heavy-chain-only IgG variants (HCAbs). In parallel, we collected animal sera, and affinity purified HCAbs with strong affinity and specificity to each antigen. The purified HCAbs were then proteolytically cleaved to generate the antigen-binding VHH fragments, which we analyzed by bottom-up Mass spectrometry (MS). Correlating peptides to the DNA database of full-length sequences using customized Llama-Magic software, we identified candidate nanobody clones. Codon-optimized genes for these clones were synthesized and cloned into a pET21-pelB E. coli periplasmic expression vector and expressed in E. coli Arctic Express (DE3) cells (Agilent); cell lysates were passed over an antigen-conjugated resin to screen for positive nanobodies.

### Creating nanobody mutants

Nanobody mutants were created by synthesizing genes with desired substitutions as gBlocks from Integrated DNA Technologies (www.idtdna.com). The gblocks (IDT) are cloned into a peT24b vector via a C-terminal BamHI and N-terminal XhoI restriction sites and correct insertions confirmed via Sanger sequencing. All mutant nanobody protein sequences are found in Supplementary Tables 5-8.

### Protein expression and purification

Protein expression and purification was performed as described previously^3^. Briefly, periplasmic expression constructs carrying the His-tagged nanobody variants were individually expressed in *E. coli* Arctic Express (DE3) cells (Agilent), periplasmic fractions released via osmotic shock, and nanobody protein purified using Ni-NTA resin (MilliporeSigma). To prepare GFP complexes for crystallization, untagged GFP was expressed in *E. coli* BL21 (DE3) cells (Agilent), which were lysed using a microfluidizer. Before elution, nanobody protein captured on Ni-NTA resin was incubated with a saturating amount of GFP in cell lysate, washed, and the GFP-nanobody complex eluted using imidazole. Eluted protein was passed over a Superdex 75 10/300 GL size-exclusion column (Cytiva) in PBS. Purity of protein preparations was analyzed via SDS-PAGE, and concentrations determined by UV absorbance or BCA assay (ThermoFisher). Samples for crystallography were concentrated by ultrafiltration when necessary.

### Protein Crystallization

Concentrations of each nanobody-GFP complex were as follows: LaG16, 10 mg mL^-1^; LaG19, 16 mg ml^-1^; LaG21, 17 mg mL^-1^; LaG24, 11 mg mL^-1^; LaG35, 13 mg mL^-1^; LaG41, 10 mg mL^-1^; LaG43, 15 mg mL^-1^. GFP-nanobody complexes at a 1:1 ratio of protein complex to reservoir buffer were crystallized via the sitting-drop vapor diffusion method where the protein solutions (0.3uL) were mixed with an equal volume of a precipitant solution and equilibrated at room temperature (294 K) against the same precipitant solution in clear tape-sealed 96-well INTELLI-plates (Art Robbins Instruments, Sunnyvale, CA) and incubated at 18 °C. Proteins were screened against the commercial screens MCSG1, MCSG2, and MCSG4 (MICROLYTIC). Crystallization was performed using either a TECAN crystallization robot (TECAN US, Research Triangle Park, NC) or a PHOENIX crystallization robot (Art Robbins Instruments). The following reservoir conditions produced crystals for subsequent data collection: LaG16-GFP complex, 0.1 M HEPES pH 7.5, 1.26 M ammonium sulphate; LaG19-GFP complex, 0.2 M lithium sulphate, 0.1 M tris pH 7.0, 1 M potassium sodium tartrate; LaG21-GFP complex, 1 M bis-tris propane pH 7.0, 1.2 M DL-malic acid pH 7.0; LaG24-GFP complex, 0.2 M potassium chloride, 20% (w/v) PEG3350; LaG35-GFP complex, 0.2 M potassium formate pH 7.3, 20% (w/v) PEG3350; LaG41-GFP, 0.2 M sodium potassium phosphate pH 6.2, 2.5 M sodium chloride; LaG43-GFP, 0.1 M sodium acetate pH 4.6, 2 M sodium formate. The crystals were harvested using cryogenic loops, cryoprotected using 20% glycerol and flash cooled in liquid nitrogen.

### Crystallographic data collection and processing

X-ray diffraction data was collected at the X29A beamline (Brookhaven National Laboratory) at a wavelength of 1.075 Å. All data were indexed, integrated and scaled with HKL3000^42^. The X-diffraction data for the LaG21-GFP complex was collected on the 31-ID beamline (Advanced Photon Source) at a wavelength of 0.9793 Å and data was indexed, integrated using MOSFLM and scaled using SCALA using the CCP4 software suit^43^. All structures were determined using molecular replacement with PHASER^44^ using as input models the GFP structure (1EMA) and the nanobody (4KRN). Each dataset underwent multiple cycles of manual editing and adjustment of the model using COOTv0.9.8.6^45^. Refinement was performed in PHENIXv1.19.2-4158^46^. The final models were validated with PROCHECK^47^, PDBePISA^32^ and images created using PyMOL (Schrodinger LLC 2015). Data collection and statistics are summarized in Supplementary Table 1.

### K_D_ determinations

K_D_’s were determined via surface plasmon resonance experiments. Measurements were either taken on a Proteon XPR36 Protein Interaction Array System (Bio-Rad) or a Biacore 8k (Cytiva) at 25 °C. Recombinant GFP was immobilized on a ProteOn GLC sensor chip as previously described ^3^. using the ProteOn Amine Coupling Kit (EDC/NHS coupling chemistry, Bio-Rad) according to the respective manufacturer’s guidelines either on a ProteOn GLC sensor chip or a Series S CM5 sensor chip. All purified nanobodies in a final buffer containing 20 mM HEPES pH 7.4, 150 mM NaCl, 0.02% Tween, were prepared in 5–8 concentrations. For experiments performed on the Proteon XPR36, protein was then injected at 50 μl/min for 120 s, followed by a dissociation time of 600 s. Residual bound proteins were removed by regenerating the chip surface using glycine pH 3, 1 M MgCl2. Data were processed and analyzed using the ProteOn Manager software. For experiments performed on the Biacore 8k, protein was injected at 60 μl/min for 120 s, followed by a dissociation time of either 1200 s or 2400s. Residual bound proteins were removed by regenerating the chip surface using 1 M MgCl2. Data were processed and analyzed using the Biacore Insight Evaluation software.

### Measurement of Melting Temperature (T_M_)

The melting temperature (T_M_) of the anti-GFP nanobody variants was measured by differential scanning fluorimetry (DSF) using a CFX96 Real Time PCR Detection System (Bio-Rad, Hercules, CA). A 96-well thin-wall PCR plate (Bio-Rad) was set up with each well containing 10-20 µM protein samples, 5 × SYPRO Orange dye (Sigma), 20 mM HEPES, 150 mM NaCl buffer (pH 7.5). The assay measured a fluorescence variation over a temperature range of 25–95°C that was increased at a rate of 0.5 °C /30 s. The excitation and emission wavelengths were 490 and 575 nm respectively. T_M_ was the transition midpoint value between the start point and the maximum point, which was calculated using the manufacturer’s software.

### Molecular modeling

Molecular dynamics (MD) simulations of solvated wildtype and mutated (L69F) LaG-21 were implemented with the openMM molecular simulation toolkit^48^, using the ff14SB amber forcefield^49^ for the protein, and a tip3p water model^50^. All simulations used a nonbonded cutoff of 10 Å, hydrogen mass of 4 amu, and timestep of 2 fs. To study the effect of temperature, we performed separate simulations at 300, 336 and 421 K. For each temperature, 8 independent copies of the system were simulated starting from slightly different initial conformations of the CDR-H3 loop (IMGT-based numbering) generated using a single iteration of the KicMover protocol, followed by 5 iterations of the FastRelax protocol in pyRosetta^44^. In each copy, the LaG21 crystal structure was first solvated using openMM’s automatic solvation routine, and then relaxed under NPT conditions at 1.013 bar pressure and 300 K temperature: first for 4 ns with 1 kcal/Å^2^ position restraints on all protein atoms, followed by further 2 ns of unrestrained simulation. The last 400 ps from the unrestrained NPT round was used to estimate the average dimensions of the cubical box required to maintain the expected water density (in the tip3p model) at 1.013 bar and 300 K. A second round of simulations was then performed under NVT conditions using the calculated box length, where the system was slowly annealed over 10 ns starting from 300 K and extending over a range of 10 exponentially distributed temperatures until the desired temperature was reached. Ultimately, an additional 100 ns of production runs under NVT conditions were carried out at the desired temperature and trajectory snapshots were recorded every 20 ps. Thus, we ran a total of 1.6 µs of production MD at each temperature, out of which the last 400 ns were used for collecting statistics. Average root mean square fluctuations of residues and backbone dihedral angles were calculated with the mdtraj Python package^45^. Overall deviation of nanobody backbone dihedral angles (*φ*, *ψ*) from the reference crystal structure (*φ*_*ref*_, *ψ*_*ref*_) was calculated for the *i^th^* residue as 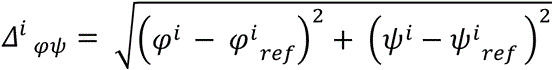 at every trajectory frame. We calculated the self (*H_i_, H_j_*) and pairwise (*H_ij_*) Gibbs entropies of the system along the backbone dihedral deviation order parameter as:

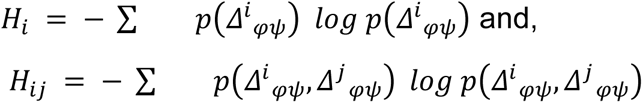

where *p(x)* is the probability distribution of the order parameter *x* (here, *Δ*_*φ ψ*_) and is calculated by histogramming the dihedral deviation data from the simulation, while the summation is carried out over all trajectory frames. Subsequently, the mutual information (*M_ij_*) between residues *i* and *j* was calculated as^34^ *M*_*ij*_ = *H*_*i*_ + *H*_*j*_ − *H*_*ij*_ and normalized to a generalized correlation coefficient between 0 and 1 as 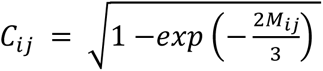.

## References

1. Muyldermans, S. Nanobodies: natural single-domain antibodies. Annu Rev Biochem 82, 775–97 (2013).

2. Mast, F.D. et al. Highly synergistic combinations of nanobodies that target SARS-CoV-2 and are resistant to escape. Elife 10(2021).

3. Fridy, P.C. et al. A robust pipeline for rapid production of versatile nanobody repertoires. Nature methods 11, 1253–1260 (2014).

4. Muyldermans, S. Applications of Nanobodies. Annu Rev Anim Biosci 9, 401–421 (2021).

5. Jovcevska, I. & Muyldermans, S. The Therapeutic Potential of Nanobodies. BioDrugs 34, 11–26 (2020).

6. Liu, M., Li, L., Jin, D. & Liu, Y. Nanobody-A versatile tool for cancer diagnosis and therapeutics. Wiley Interdiscip Rev Nanomed Nanobiotechnol 13, e1697 (2021).

7. Li, C. et al. Broad neutralization of SARS-CoV-2 variants by an inhalable bispecific single-domain antibody. Cell 185, 1389–1401 e18 (2022).

8. Pymm, P. et al. Nanobody cocktails potently neutralize SARS-CoV-2 D614G N501Y variant and protect mice. Proc Natl Acad Sci U S A 118(2021).

9. Chi, X. et al. An ultrapotent RBD-targeted biparatopic nanobody neutralizes broad SARS-CoV-2 variants. Signal Transduct Target Ther 7, 44 (2022).

10. Wu, X. et al. A potent bispecific nanobody protects hACE2 mice against SARS-CoV-2 infection via intranasal administration. Cell Rep 37, 109869 (2021).

11. Romao, E. et al. Identification of Useful Nanobodies by Phage Display of Immune Single Domain Libraries Derived from Camelid Heavy Chain Antibodies. Curr Pharm Des 22, 6500–6518 (2016).

12. Muyldermans, S., et al. Camelid immunoglobulins and nanobody technology. Veterinary immunology and immunopathology 128, 178–183 (2009).

13. Arbabi Ghahroudi, M., Desmyter, A., Wyns, L., Hamers, R. & Muyldermans, S. Selection and identification of single domain antibody fragments from camel heavy-chain antibodies. FEBS Lett 414, 521–6 (1997).

14. Vincke, C. et al. General strategy to humanize a camelid single-domain antibody and identification of a universal humanized nanobody scaffold. Journal of Biological Chemistry 284, 3273–3284 (2009).

15. Kunz, P. et al. The structural basis of nanobody unfolding reversibility and thermoresistance. Sci Rep 8, 7934 (2018).

16. Govaert, J. et al. Dual beneficial effect of interloop disulfide bond for single domain antibody fragments. J Biol Chem 287, 1970–9 (2012).

17. Goldman, E.R., Liu, J.L., Zabetakis, D. & Anderson, G.P. Enhancing Stability of Camelid and Shark Single Domain Antibodies: An Overview. Front Immunol 8, 865 (2017).

18. Kunz, P., Ortale, A., Mucke, N., Zinner, K. & Hoheisel, J.D. Nanobody stability engineering by employing the DeltaTm shift; a comparison with apparent rate constants of heat-induced aggregation. Protein Eng Des Sel 32, 241–249 (2019).

19. Dingus, J.G., Tang, J.C.Y., Amamoto, R., Wallick, G.K. & Cepko, C.L. A general approach for stabilizing nanobodies for intracellular expression. Elife 11(2022).

20. Saerens, D., Conrath, K., Govaert, J. & Muyldermans, S. Disulfide bond introduction for general stabilization of immunoglobulin heavy-chain variable domains. Journal of molecular biology 377, 478–488 (2008).

21. Kunz, P. et al. Exploiting sequence and stability information for directing nanobody stability engineering. Biochimica et Biophysica Acta (BBA)-General Subjects 1861, 2196–2205 (2017).

22. De Genst, E. et al. Molecular basis for the preferential cleft recognition by dromedary heavy-chain antibodies. Proc Natl Acad Sci U S A 103, 4586–91 (2006).

23. Mitchell, L.S. & Colwell, L.J. Comparative analysis of nanobody sequence and structure data. Proteins 86, 697–706 (2018).

24. Zimmermann, I. et al. Synthetic single domain antibodies for the conformational trapping of membrane proteins. Elife 7(2018).

25. Castell, S.E., Jordan, S.L. & Halford, S.E. Site-specific recombination and topoisomerization by Tn21 resolvase: role of metal ions. Nucleic Acids Res 14, 7213–26 (1986).

26. Zavrtanik, U., Lukan, J., Loris, R., Lah, J. & Hadzi, S. Structural Basis of Epitope Recognition by Heavy-Chain Camelid Antibodies. J Mol Biol 430, 4369–4386 (2018).

27. Endo, H. et al. Role of the giant panda’s ’pseudo-thumb’. Nature 397, 309–10 (1999).

28. Kirchhofer, A. et al. Modulation of protein properties in living cells using nanobodies. Nature structural & molecular biology 17, 133 (2010).

29. Kazemi-Lomedasht, F., Muyldermans, S., Habibi-Anbouhi, M. & Behdani, M. Design of a humanized anti vascular endothelial growth factor nanobody and evaluation of its in vitro function. Iran J Basic Med Sci 21, 260–266 (2018).

30. Moutel, S. et al. NaLi-H1: A universal synthetic library of humanized nanobodies providing highly functional antibodies and intrabodies. Elife 5(2016).

31. Soler, M.A. et al. Effect of Humanizing Mutations on the Stability of the Llama Single-Domain Variable Region. Biomolecules 11(2021).

32. Krissinel, E. & Henrick, K. Inference of macromolecular assemblies from crystalline state. Journal of molecular biology 372, 774–797 (2007).

33. Bond, C.J., Marsters, J.C. & Sidhu, S.S. Contributions of CDR3 to V H H domain stability and the design of monobody scaffolds for naive antibody libraries. J Mol Biol 332, 643–55 (2003).

34. Lange, O.F. & Grubmuller, H. Generalized correlation for biomolecular dynamics. Proteins 62, 1053–61 (2006).

35. Martinez-Delgado, G. Inhaled nanobodies against COVID-19. Nat Rev Immunol 20, 593 (2020).

36. Yang, E.Y. & Shah, K. Nanobodies: Next Generation of Cancer Diagnostics and Therapeutics. Front Oncol 10, 1182 (2020).

37. Bao, C. et al. The Application of Nanobody in CAR-T Therapy. Biomolecules 11(2021).

38. Marable, J. et al. Nanobody-based CTLA4 inhibitors for immune checkpoint blockade therapy of canine cancer patients. Sci Rep 11, 20763 (2021).

39. Deken, M.M. et al. Nanobody-targeted photodynamic therapy induces significant tumor regression of trastuzumab-resistant HER2-positive breast cancer, after a single treatment session. J Control Release 323, 269–281 (2020).

40. Messer, A. & Butler, D.C. Optimizing intracellular antibodies (intrabodies/nanobodies) to treat neurodegenerative disorders. Neurobiol Dis 134, 104619 (2020).

41. Wouters, Y. et al. VHHs as tools for therapeutic protein delivery to the central nervous system. Fluids Barriers CNS 19, 79 (2022).

42. Otwinowski, Z. & Minor, W. Processing of X-ray diffraction data. Methods enzymol 276, 307–326 (1997).

43. Winn, M.D. et al. Overview of the CCP4 suite and current developments. Acta Crystallographica Section D: Biological Crystallography 67, 235–242 (2011).

44. McCoy, A.J., et al. Phaser crystallographic software. Journal of applied crystallography 40, 658–674 (2007).

45. Emsley, P. & Cowtan, K. Coot: model-building tools for molecular graphics. Acta Crystallographica Section D: Biological Crystallography 60, 2126–2132 (2004).

46. Liebschner, D. et al. Macromolecular structure determination using X-rays, neutrons and electrons: recent developments in Phenix. Acta Crystallogr D Struct Biol 75, 861–877 (2019).

47. Laskowski, R.A., MacArthur, M.W., Moss, D.S. & Thornton, J.M. PROCHECK: a program to check the stereochemical quality of protein structures. Journal of applied crystallography 26, 283–291 (1993).

48. Eastman, P. et al. OpenMM 7: Rapid development of high performance algorithms for molecular dynamics. PLoS Comput Biol 13, e1005659 (2017).

49. Maier, J.A. et al. ff14SB: Improving the Accuracy of Protein Side Chain and Backbone Parameters from ff99SB. J Chem Theory Comput 11, 3696–713 (2015).

50. Jorgensen, W.L., Chandrasekhar, J., Madura, J.D., Impey, R.W. & Klein, M.L. Comparison of simple potential functions for simulating liquid water. The Journal of chemical physics 79, 926–935 (1983).

