## Supplementary Figures and Tables for "Unique Binding and Stabilization Mechanisms Employed By and Engineered Into Nanobodies"

### Supplemental Figures and Tables

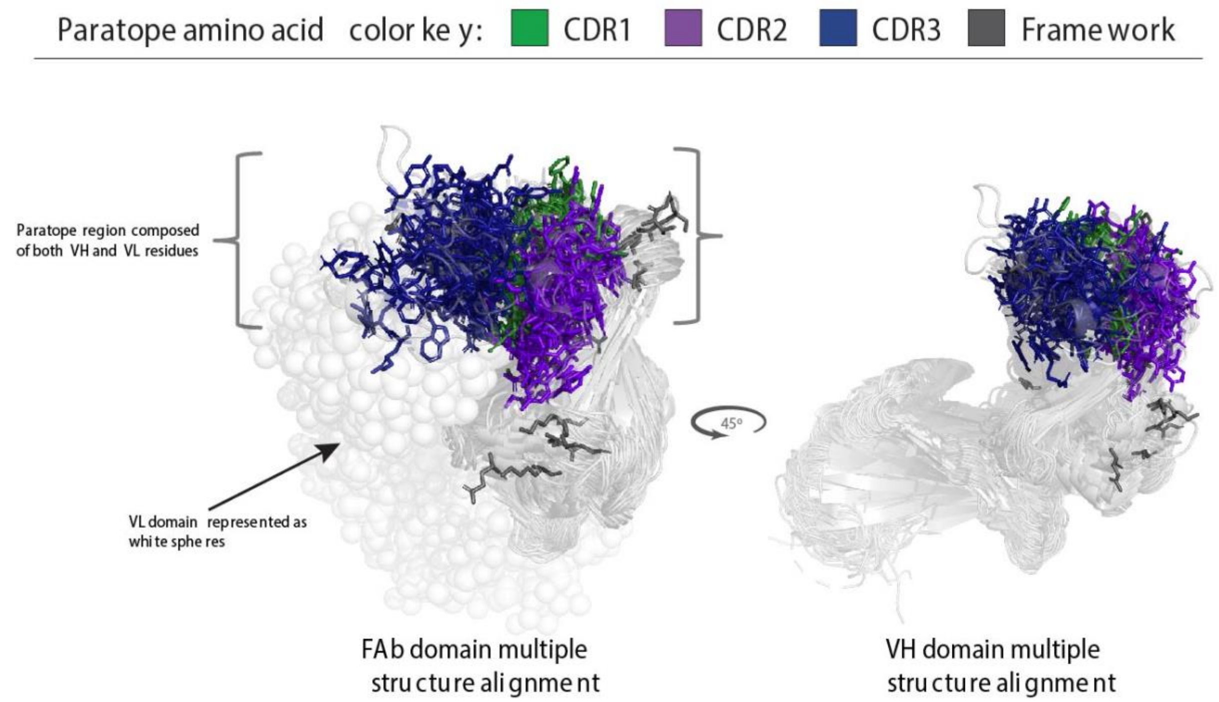


**Supplementary Figure 1. Multiple structure alignment of 87 FAb structures in complex with their antigen (antigen not shown).** The crystal structures were aligned in PyMOL (Schrodinger LLC 2015) where the VH domain are highlighted. The paratope (binding site) is represented in stick form and color coded using the color key. The PDBs used in the multiple structure alignment are found in Supplementary Table 3.

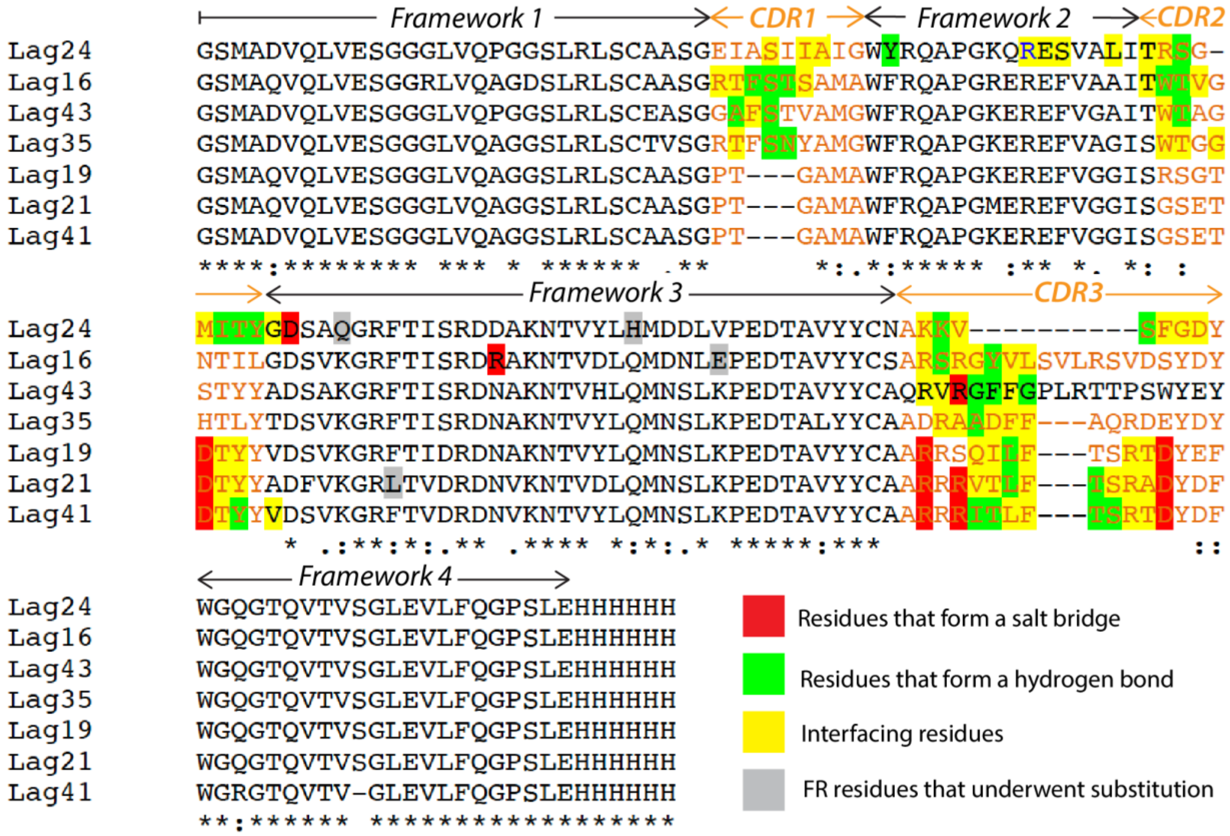


**Supplementary Figure 2. Sequence alignment of the seven LaG nanobodies whose structures were solved in this study.** The multiple sequence alignment was generated using Clustal Omega[48]. Assignment of framework and CDR regions is according to IMGT numbering. Visual inspection using PyMOL coupled to PBDePISA[49] analysis was used to determine residues at the protein-protein interface of each LaG-GFP complex. Interface residues are colored according to their role at the interface.

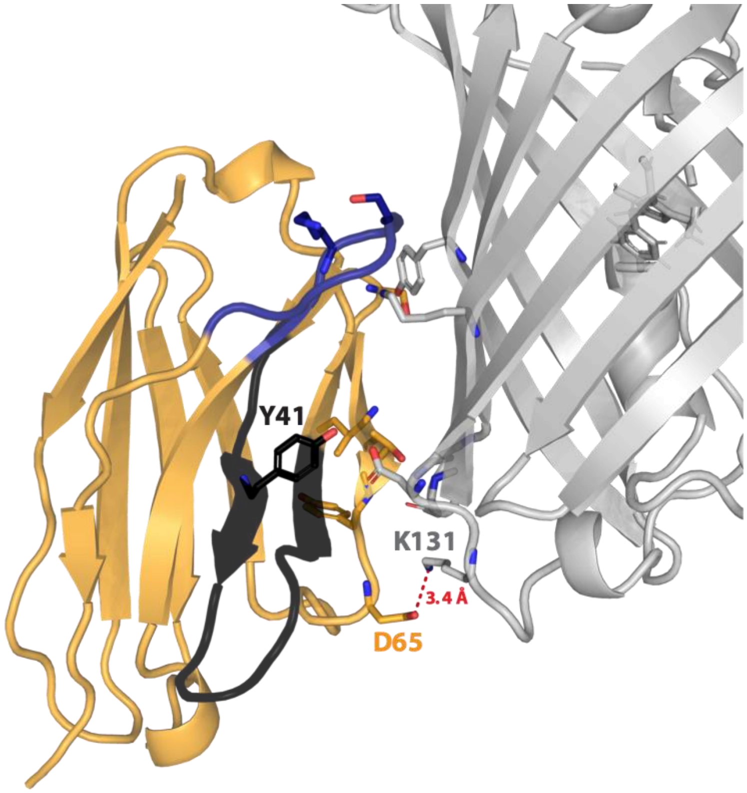


**Supplementary Figure 3. Close up of the LaG24-GFP interface.** The side-on interactor LaG24 has a shorter CDR3 (*blue*), which opens up its FW2 region (*black*) to interact with GFP. Additionally, the only electrostatic interaction formed with GFP is *via* Asp65 at the periphery of the paratope, which may serve to secure the interaction interface.

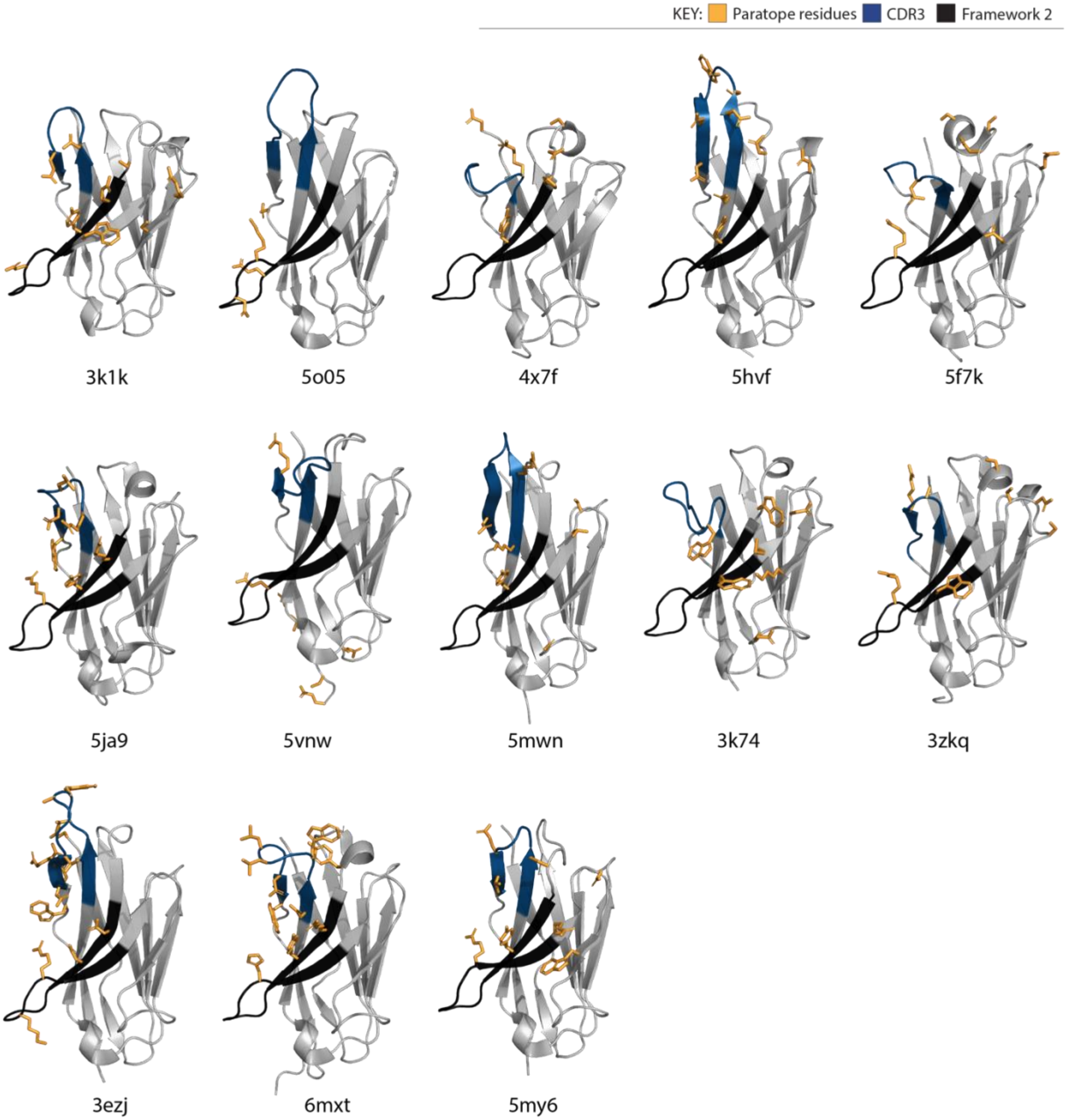


**Supplementary Figure 4.** Examples of 13 nanobodies in the PDB that interact side-on with their antigen, promoted by their shorter CDR3.

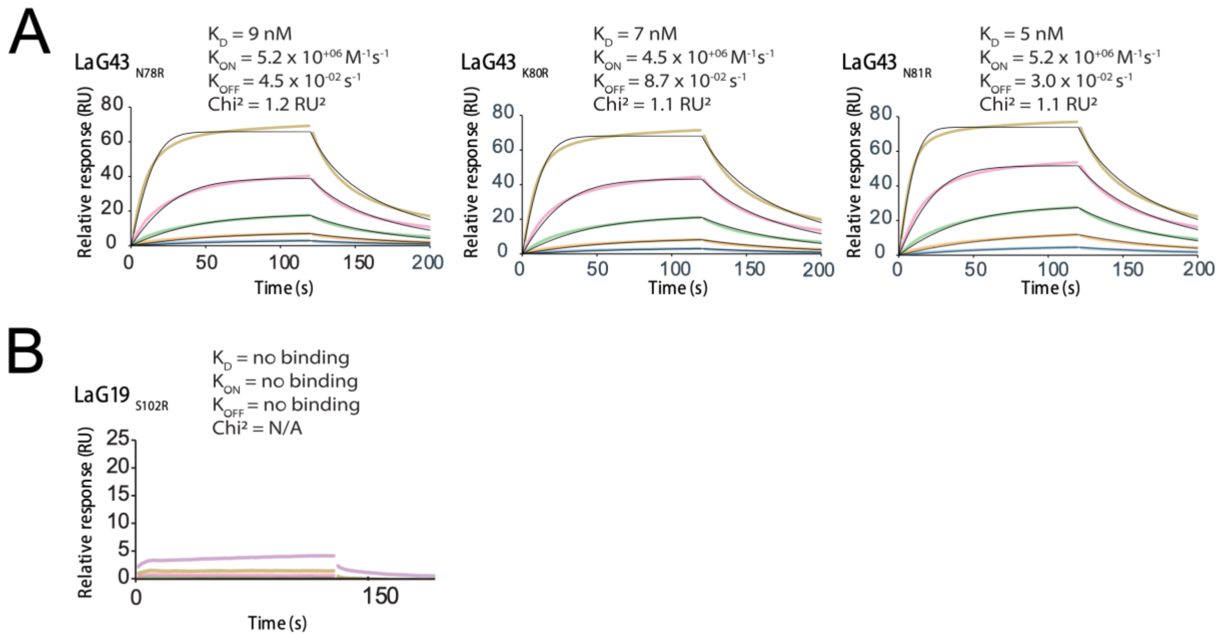


**Supplementary Figure 5. SPR results of reengineered nanobody paratopes against GFP point mutants. A)** SPR results of the three LaG43 point mutants against the $GFP_{E17A}$ point variant that removes the glutamic acid which we hypothesize interacts with the point variants with a single arginine introduced at different positions along the length of the 4$^{th}$ loop on LaG43. As predicted, the direct **B)** SPR result of $LaG19_{S102R}$ against the $GFP_{E32A}$ point variant that removes the glutamic acid which we hypothesize interacts with the introduced arginine.

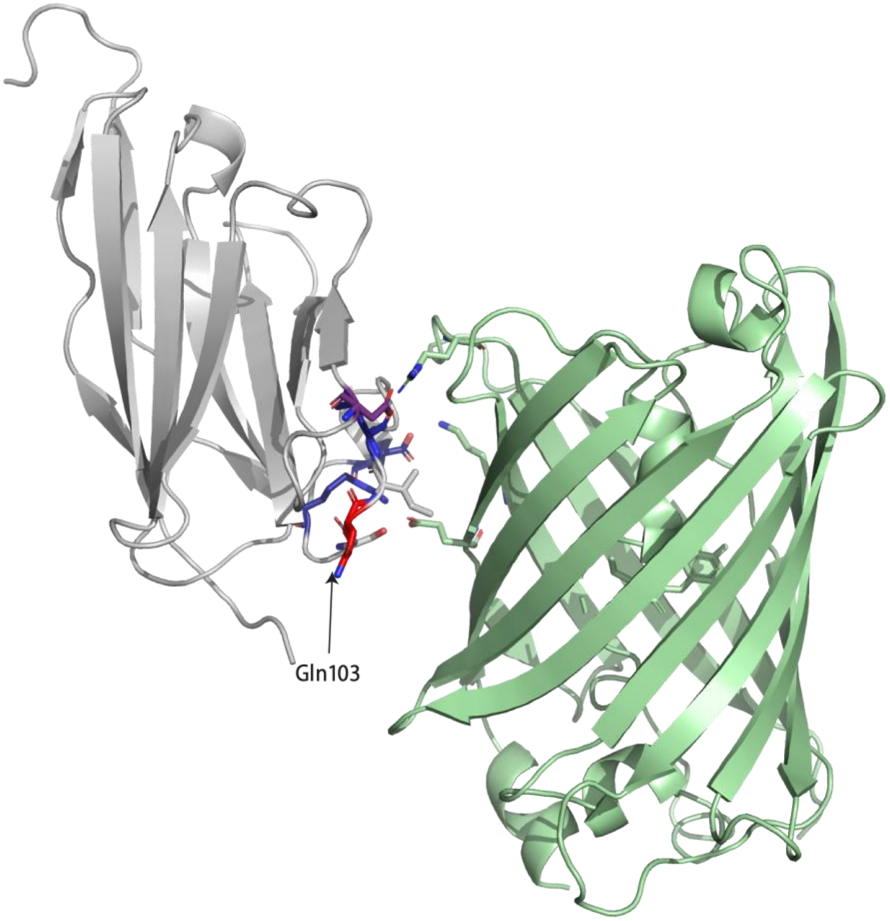


**Supplementary Figure 6. LaG19-GFP complex.** The paratope residues of LaG19 (*grey*) and GFP (*green*) are represented as sticks. The exception for LaG19 is the two stick residues colored in *grey*, which flank Gln103, the residue mutated when probing paratope reengineering - relating to Figure 3 in the main text.

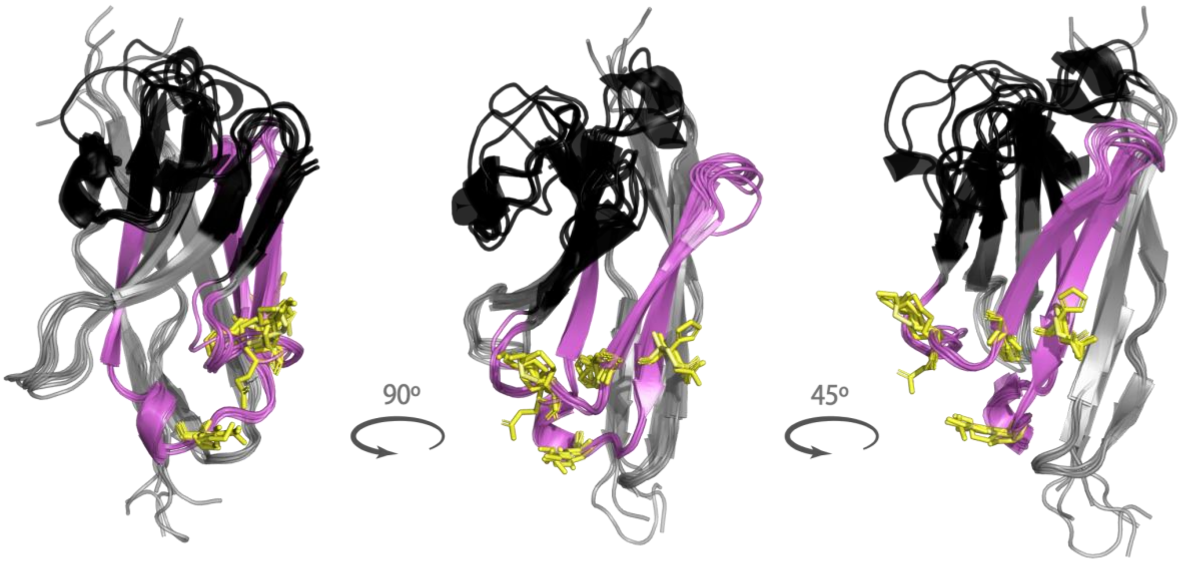


**Supplementary Figure 7. The location of the FW3 residues that enable the fine-tuning of nanobody affinity and/or stability.** In yellow is the location of key regions identified in this study on FW3 (*pink*), shown to influence nanobody affinity and/or stability. The location of these residues is on the opposite face of the nanobody to the CDR loops (*black*).

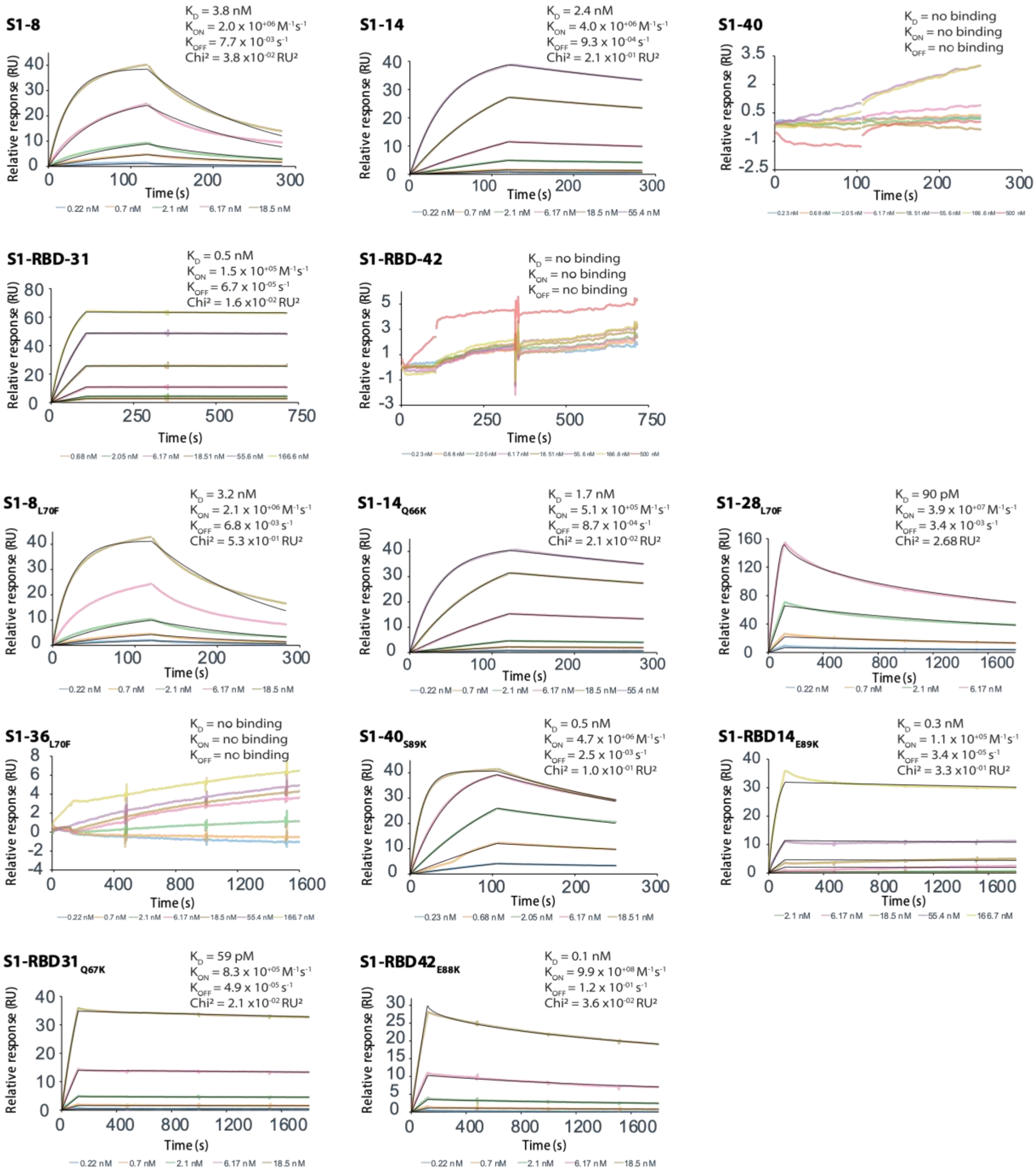


**Supplementary Figure 8. SPR sensograms relating to Table 1 (main text).**

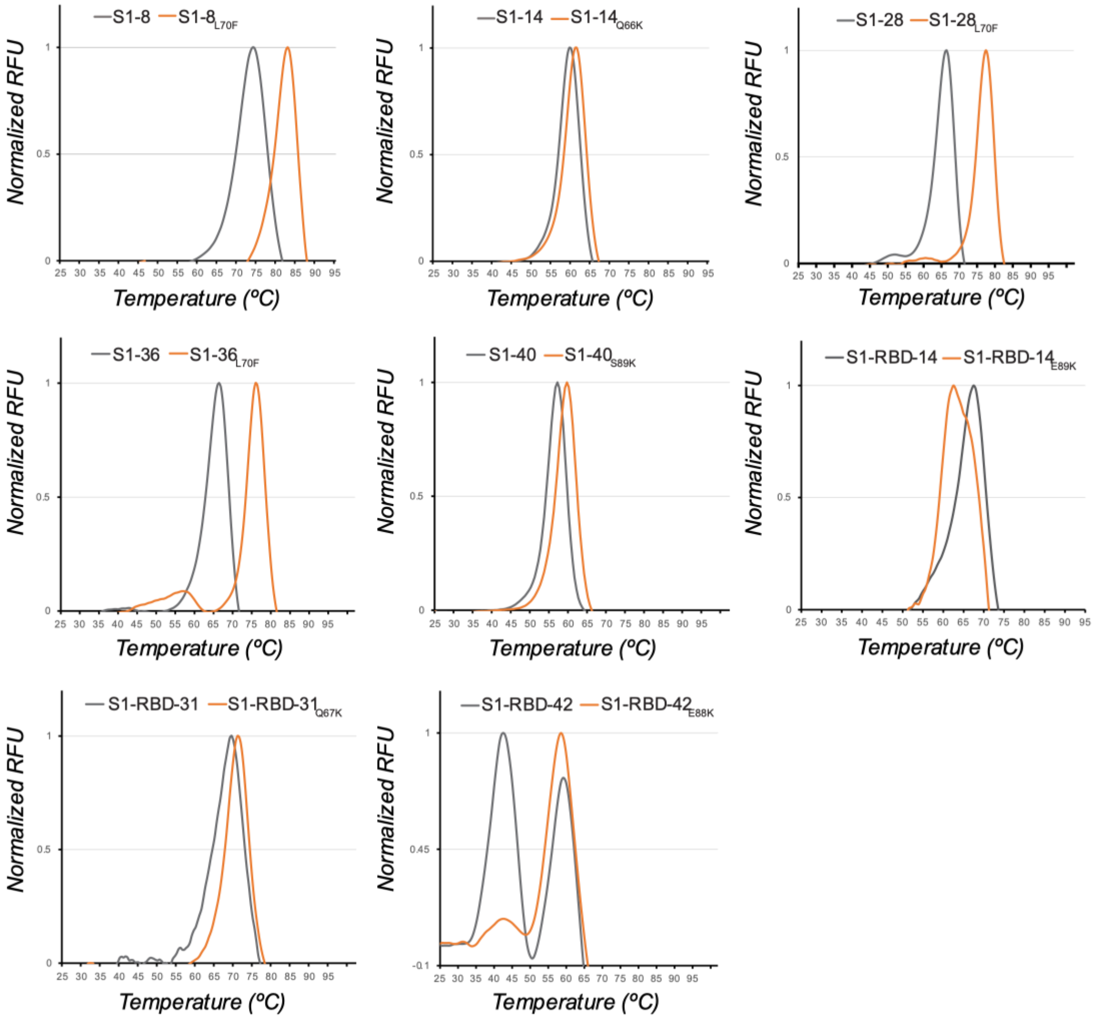


**Supplementary Figure 9. DSF results of nanobodies from Table 1.** Thermal denaturation profiles of anti-SARS-CoV-2 S1 nanobodies (*grey*) compared to their corresponding point variants (*orange*).

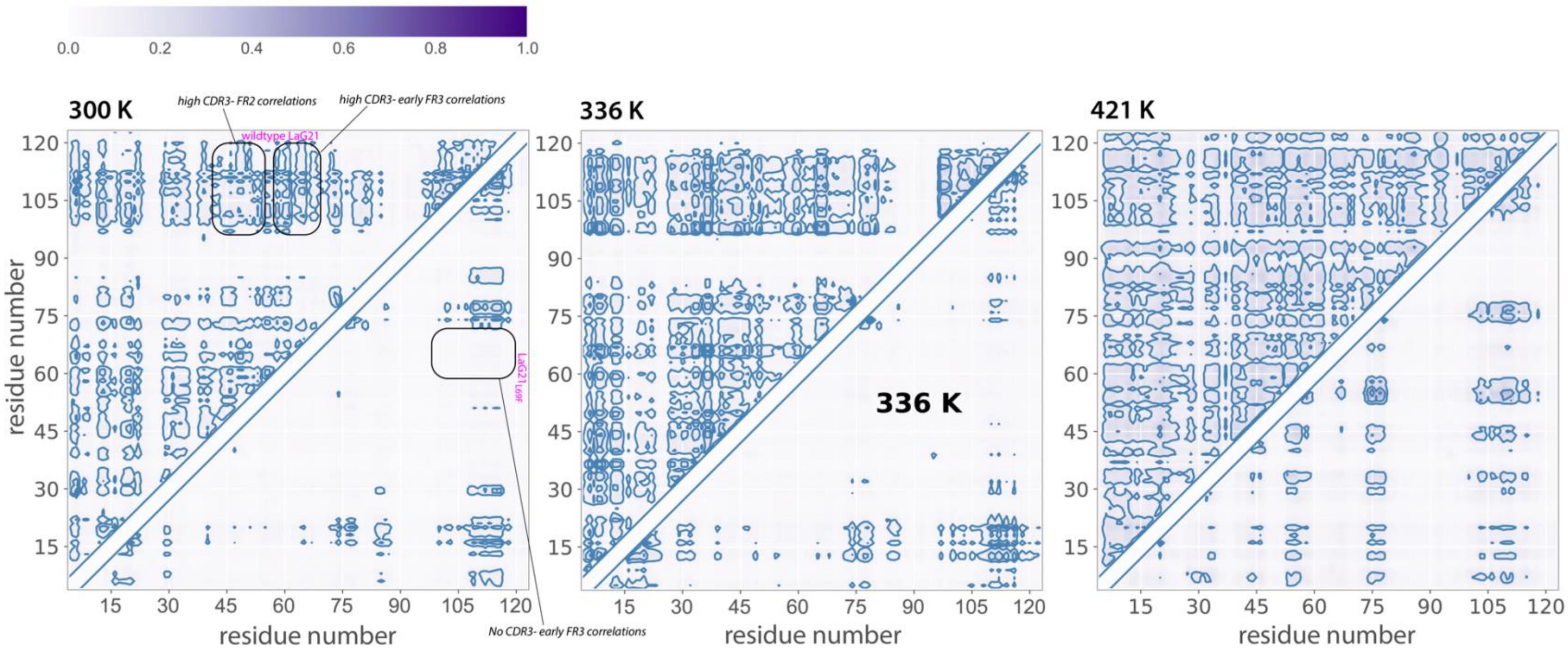


**Supplementary Figure 10. Mutual Information (MI) analysis of LaG21 and LaG21$_{L69F}$.** Normalized mutual information between backbone dihedral angles for residue pairs at different temperatures for LaG21 above the diagonal and LaG21$_{L69F}$ below the diagonal. Presence of the Leu69Phe mutation likely reduces correlated motion with both CDR2 and CDR3 (boxed in black) leading to nearly zero mutual information between the CDR3 and FRs 2-4 (section along Y axis), On the other hand, in the WT we observe very high "communication" between structurally distant domain-pairs like CDR3-FR3 and even CDR3-FR2 (section along X axis).

**Supplementary Table 1. Data collection and refinement statistics**

| **Structure:**<br>**PDB ID:** | **LaG16**<br>**8SFS** | **LaG43**<br>**8SLC** | **LaG24**<br>**8G0I** | **LaG19**<br>**8SFV** | **LaG21**<br>**8SFX** | **LaG41**<br>**8SG3** | **LaG35**<br>**8SFZ** |
|---|---|---|---|---|---|---|---|
| **Data collection** | | | | | | | |
| Space group | $P3_121$ | P63 | $P\ 2_12_12$ | $I4_122$ | I4 | $I4_1$ | $P22_12_1$ |
| Cell dimensions | | | | | | | |
| *a*, *b*, *c* (Å) | 131.77, 131.77, 153.58 | 149.43, 149.43, 127.01 | 82.1, 86.7, 52.1 | 111.25, 111.25, 193.8 | 111.12, 111.12, 194.20 | 108.63, 108.63, 198.74 | 69.65, 101.73, 184.48 |
| α, β, γ (°) | 90, 90, 120 | 90, 90, 120 | 90, 90, 90 | 90, 90, 90 | 90, 90, 90 | 90, 90, 90 | 90, 90, 90 |
| Resolution (Å) | 50.0 – 2.20 (2.24 – 2.20) | 50.0-2.70 (2.75-2.70) | 50.0 – 2.20 (2.26 – 2.20) | 41.25 – 1.80 (1.83 – 1.80) | 41.3 – 1.95 (2.00 – 1.95) | 50 – 3.0 (3.05 – 3.0) | 50.0 – 1.9 (1.93 – 1.9) |
| $R_{sym}$ or $R_{merge}$ | 0.107(0.962) | 0.117(0.941) | 0.089(0.726) | 0.083(0.908) | 0.130(1.403) | 0.212(0.833) | 0.086(0.314) |
| $I / \sigma I$ | 22.6(1.67) | 20.4(1.13) | 22.4(2.61) | 29.5(2.83) | 11.1(2.0) | 10.3(1.78) | 14.8(1.51) |
| Completeness (%) | 95.9(69.0) | 99.9(98.8) | 99.5(93.4) | 99.6(93.1) | 94.4(81.88) | 84.6(94.0) | 91.2(62.8) |
| Redundancy | 10.4(8.4) | 4.4(4.2) | 10.0(6.2) | 14.3(14.5) | 15.1(12.7) | 3.7(3.9) | 3.8(3.1) |
| **Refinement** | | | | | | | |
| Resolution (Å) | 46.7-2.37 | 48.39 – 2.97 | 44.65 – 2.20 | 41.25 – 1.83 | 36.67 – 1.95 | 47.19 – 3.11 | 46.10 – 1.90 |
| No. reflections | 62729 | 33003 | 18349 | 51900 | 80508 | 17265 | 103243 |
| $R_{work}$ / $R_{free}$ | 0.1904/0.2170 | 0.2184/0.2399 | 0.1811/0.2413 | 0.1827/0.2059 | 0.2101/0.2275 | 0.2746/0.3044 | 0.2067/0.2312 |
| No. atoms | | | | | | | |
| Protein | 5563 | 5562 | 2632 | 2793 | 5402 | 2754 | 8284 |
| Ligand/ion | 275 | 118 | 38 | 108 | 212 | 31 | 96 |
| Water | 193 | 9 | 66 | 170 | 146 | 6 | 406 |
| *B*-factors | | | | | | | |
| Protein | 56.96 | 69.71 | 44.78 | 45.13 | 50.84 | 77.80 | 46.59 |
| Ligand/ion | 76.89 | 68.07 | 44.83 | 72.76 | 70.23 | 63.69 | 42.79 |
| Water | 54.25 | 51.65 | 43.40 | 46.95 | 46.00 | 51.5 | 42.93 |
| R.m.s. deviations | | | | | | | |
| Bond lengths (Å) | 0.008 | 0.006 | 0.013 | 0.011 | 0.012 | 0.004 | 0.012 |
| Bond angles (°) | 0.95 | 0.94 | 1.23 | 1.13 | 1.11 | 0.79 | 1.05 |

Values in parentheses are for highest-resolution shell.

**Supplementary Table 2. 95 PDB IDs of nanobody-antigen complexes used to generate Fig. 1D and Fig. 4A.**

| Interaction mode | pdb_code | species | chain_id |
|---|---|---|---|
| **Head-on** | **1BZQ** | Camelus dromedarius | N |
| | **4CDG** | Lama glama | D |
| | **4GFT** | Lama glama | B |
| | **4KRM** | Lama glama | H |
| | **4KRO** | Lama glama | B |
| | **4KRP** | Lama glama | B |
| | **4P2C** | Lama glama | G |
| | **6H02** | Lama glama | B |
| | **6B20** | Lama glama | E |
| | **6EY6** | Lama glama | M |
| | **6EQI** | Lama glama | B |
| | **4S10** | Lama glama | A |
| | **5IP4** | Lama glama | B |
| | **5M94** | Lama glama | D |
| | **5UK4** | Vicugna pacos | r |
| | **5OVW** | Lama glama | L |
| | **5O0W** | Vicugna pacos | E |
| **Side-on** | **4W6W** | Lama glama | B |
| | **3K1K** | Camelus dromedarius | C |
| | **3ZKQ** | Lama glama | D |
| | **4C58** | Lama glama | B |
| | **4KML** | Lama glama | B |
| | **4LGP** | Vicugna pacos | D |
| | **4QO1** | Lama glama | A |
| | **4X7F** | Vicugna pacos | C |
| | **4Y8D** | Lama glama | D |
| | **5C1M** | Lama glama | B |
| | **5C3L** | Camelus dromedarius | D |
| | **5DA0** | Lama glama | B |
| | **5F7K** | Lama glama | D |
| | **6MXT** | Lama glama | N |
| | **5USF** | Lama glama | C |
| | **5VNW** | synthetic | C |
| | **5JA9** | Lama glama | A |
| | **5M2M** | Lama glama | H |
| | **5M30** | Vicugna pacos | E |
| | **5MY6** | Camelus dromedarius | B |
| | **5NBD** | Lama glama | C |
| | **5O03** | Lama glama | C |
| | **5O05** | Vicugna pacos | C |
| | **5OMM** | Vicugna pacos | C |

| | | | |
|---|---|---|---|
| | **5OMN** | Vicugna pacos | C |
| | **5TOK** | Lama glama | E |
| | **2X6M** | Camelus dromedarius | A |
| | **3K74** | Lama glama | B |
| | **4AQ1** | Lama glama | B |
| | **4I0C** | Camelus dromedarius | C |
| | **5MWN** | Lama glama | F |
| | **5TOJ** | Lama glama | E |
| **Side-on CDR3** | **1JTO** | Camelus dromedarius | A |
| | **3CFI** | Lama glama | C |
| | **3EBA** | Camelus dromedarius | A |
| | **3EZJ** | Lama glama | B |
| | **3G9A** | Camelus dromedarius | B |
| | **3STB** | Lama glama | B |
| | **4C57** | Lama glama | D |
| | **4EIZ** | Lama glama | C |
| | **4FHB** | Lama glama | D |
| | **4HEM** | Lama glama | G |
| | **4HEP** | Lama glama | G |
| | **4MQS** | Lama glama | B |
| | **4N9O** | Lama glama | B |
| | **4OCM** | Lama glama | C |
| | **4OCN** | Lama glama | F |
| | **4W6X** | Lama glama | B |
| | **4WEM** | Lama glama | B |
| | **4X7C** | Vicugna pacos | D |
| | **4X7D** | Lama glama | C |
| | **4X7E** | Vicugna pacos | D |
| | **4XT1** | Vicugna pacos | C |
| | **5C2U** | Camelus dromedarius | B |
| | **5E7F** | Camelus dromedarius | C |
| | **5F7L** | Lama glama | B |
| | **5NBL** | Vicugna pacos | F |
| | **6C9W** | Lama glama | B |
| | **6FE4** | Vicugna pacos | J |
| | **6EHG** | Lama glama | C |
| | **6EY0** | Lama glama | E |
| | **5VAK** | Lama glama | B |
| | **5VAN** | Lama glama | B |
| | **5VAQ** | Lama glama | B |
| | **5MP2** | Lama glama | D |
| | **5O8F** | Lama glama | L |
| | **5HVF** | Vicugna pacos | B |

| | | | |
|---|---|---|---|
| | **5HVG** | Vicugna pacos | B |
| | **5IMK** | Camelidiae | B |
| | **5JA8** | Lama glama | H |
| | **5JDS** | Camelus bactrianus | B |
| | **5JQH** | Lama glama | D |
| | **5LHN** | Vicugna pacos | B |
| | **5LHR** | Vicugna pacos | B |
| | **5MJE** | Lama glama | B |
| | **5O02** | Vicugna pacos | C |
| | **5OJM** | Lama glama | L |
| | **5TJW** | Vicugna pacos | K |

**Supplementary Table 3. 87 PDB IDs of FAb-antigen complexes used to generate Supplementary Fig. 1 and Fig. 4B**

| pdb_code | chain_id |
|---|---|
| 1CU4 | H |
| 5UOE | H |
| 5BJZ | C |
| 6AYZ | B |
| 5NJD | R |
| 5W6G | H |
| 5W23 | H |
| 5OB5 | H |
| 5GUX | H |
| 5MU2 | G |
| 5VPG | D |
| 1BJ1 | H |
| 5MY4 | B |
| 5G64 | I |
| 5VIC | H |
| 5VCN | D |
| 5SY8 | H |
| 5U3K | H |
| 5MEV | H |
| 5H35 | F |
| 5KTE | H |
| 5SX4 | J |
| 5EU7 | E |
| 5HDQ | H |
| 5GIR | A |
| 5 E 94 | D |
| 5J9P | A |
| 5E1A | A |
| 5DHZ | H |
| 5FUO | H |
| 5HHV | H |
| 5B3J | E |
| 5EPM | A |
| 5FGB | C |
| 5DFV | C |

| | |
|---|---|
| 5E8D | H |
| 5BO1 | H |
| 4ZPT | H |
| 5C7X | H |
| 4YX2 | H |
| 4WV1 | E |
| 4ZS6 | H |
| 1NDM | B |
| 1FJ1 | D |
| 1JPS | H |
| 3THM | H |
| 6FAX | H |
| 2HFG | H |
| 2OSL | A |
| 3FMG | H |
| 4QTI | H |
| 4ZFF | H |
| 1LK3 | H |
| 2ZCL | H |
| 2ARJ | B |
| 1SY6 | H |
| 2QQN | H |
| 3R08 | H |
| 2B2X | H |
| 5NUZ | H |
| 5TZ2 | H |
| 5VL3 | A |
| 4AG4 | H |
| 3UJI | H |
| 4EDW | H |
| 4JO1 | H |
| 4F3F | B |
| 5O14 | H |
| 4P9H | H |
| 4Q6I | B |
| 3PGF | H |
| 3KS0 | K |
| 4K2U | I |
| 2BRR | H |
| 3WIH | H |
| 3IU3 | H |
| 4QEX | H |
| 2H9G | B |
| 4I2X | D |
| 1EJO | H |
| 3NH7 | J |
| 3BGF | H |
| 3EOA | H |
| 1V7M | I |

#### Supplementary Table 4: List of direct contacts between nanobodies and GFP

**Hydrogen bonds**

| LaG16 | Dist. (Å) | GFP | LaG43 | Dist. (Å) | GFP | LaG24 | Dist. (Å) | GFP | LaG35 | Dist. (Å) | GFP |
|---|---|---|---|---|---|---|---|---|---|---|---|
| Phe33 (O) | 2.67 | Arg122 (NH2) | Ala28 (O) | 2.90 | Arg109 (NH2) | Tyr41 (HH) | 1.99 | Asp129 (OD2) | Ser31 (O) | 2.26 | Asn198 (HD22) |
| Ser34 (O) | 2.93 | Lys113 (NZ) | Ser30 (H) | 2.31 | Glu111 (OE2) | Ser58 (H) | 2.11 | Tyr182 (OH) | Asn32 (OD1) | 2.03 | Asn198 (H) |
| Thr35 (O) | 2.75 | Arg122 (NH2) | Ser30 (OG) | 3.59 | Glu111 (OE1) | Tyr63 (H) | 2.17 | Asp102 (OD1) | Ala103 (O) | 1.97 | Lys162 (HZ2) |
| Thr58 (OG1) | 3.07 | Arg122 (NH1) | Ser30 (O) | 3.00 | Lys113 (NZ) | Ile61 (O) | 2.36 | Lys101 (HZ2) | Asn32 (HD21) | 2.23 | Asp197 (OD1) |
| Ser104 (O) | 3.04 | Lys113 (NZ) | Thr54 (OG1) | 3.00 | Arg122 (NH2) | Thr62 (OG1) | 2.02 | Asp102 (H) | | | |
| Tyr107 (O) | 2.97 | Gly116 (N) | Arg101 (NH1) | 3.55 | Asp190 (OD1) | Lys103 (O) | 2.18 | Lys107 (HZ3) | | | |
| Arg78 (NH2) | 2.75 | Glu17 (OE1) | Gly102 (H) | 2.11 | Glu90 (O) | Ser105 (O) | 2.41 | Lys107 (HZ3) | | | |
| Arg78 (NH1) | 3.18 | Glu17 (OE2) | Phe103 (H) | 2.07 | Phe114 (O) | | | | | | |
| Tyr107 (N) | 3.89 | Pro89 (O) | Gly105 (H) | 2.07 | Gly116 (O) | | | | | | |
| Ser34 (OG) | 2.58 | Glu111 (OE2) | | | | | | | | | |
| Tyr107 (N) | 2.91 | Phe114 (O) | | | | | | | | | |
| Trp 57 (N) | 3.22 | Glu115 (OE1) | | | | | | | | | |
| Thr58 (N) | 3.06 | Glu115 (OE1) | | | | | | | | | |
| Tyr107 (OH) | 3.76 | Glu115 (OE1) | | | | | | | | | |
| Thr58 (OG1) | 2.64 | Glu115 | | | | | | | | | |

**Salt Bridges**

| LaG16 | Dist. (Å) | GFP | LaG43 | Dist. (Å) | GFP | LaG24 | Dist. (Å) | GFP |
|---|---|---|---|---|---|---|---|---|
| Arg78 (NH2) | 2.75 | Glu 17 (OE1) | Arg101 (NH1) | 3.55 | Asp190 (OD1) | Asp65 (OD1) | 3.41 | Lys131 (NZ) |
| Arg78 (NH1) | 3.18 | Glu17 (OE2) | Arg101 (NH2) | 3.78 | Asp190 (OD1) | | | |
| Arg78 (NH2) | 3.22 | Glu 17 (OE1) | | | | | | |

**Hydrogen bonds**

| LaG41 | Dist. (Å) | GFP | LaG21 | Dist. (Å) | GFP | LaG19 | Dist. (Å) | GFP |
|---|---|---|---|---|---|---|---|---|
| Asp111 (OD2) | 2.85 | Lys45 (HZ2) | Arg100 (HH22) | 1.56 | Glu32 (OE2) | Asp111 (OD2) | 1.80 | Lys45 (HZ3) |
| Ser108 (OG) | 3.29 | Glu213 (N) | Arg102 (NH1) | 3.11 | Glu32 (OE2) | Asp58 (OD2) | 1.85 | Arg215 (HH12) |
| Thr107 (OG1) | 2.33 | Lys214 (N) | Arg102 (NH2) | 2.68 | Glu32 (O) | Leu105 (O) | 2.23 | Arg215 (HH11) |
| Thr107 (OG1) | 3.59 | Arg215 (N) | Thr107 (OG1) | 3.74 | Asn212 (O) | Asp58 (OD1) | 1.96 | Arg215 (HH22) |
| Asp58 (OD2) | 1.74 | Arg215 (HH12) | Thr107 (H) | 2.07 | Glu213 (OE1) | | | |
| Arg100 (NH2) | 2.77 | Glu32 (OE1) | Asp111 (OD2) | 1.84 | Lys45 (HZ1) | | | |
| Thr104 (N) | 2.73 | Glu32 (OE2) | Asp58 (OD2) | 1.74 | Arg215 (HH12) | | | |
| Ile103 (N) | 3.38 | Glu32 (OE2) | Leu105 (O) | 2.20 | Arg215 (HH11) | | | |
| Arg100 (NH1) | 2.70 | Glue32 (OE2) | Asp58 (OD1) | 1.99 | Arg215 (HH22) | | | |
| Ser108 (OG) | 2.42 | Asp210 (OD2) | | | | | | |
| Thr107 (OG1) | 3.59 | Asn212 (OD1) | | | | | | |
| Ser108 (N) | 3.51 | Glu213 (OE1) | | | | | | |
| Thr107 (N) | 2.85 | Glu213 (OE1) | | | | | | |
| Ser108 (OG) | 3.35 | Glu213 (OE1) | | | | | | |
| Thr107 (N) | 2.90 | Glu213 (OE2) | | | | | | |
| Tyr60 (OH) | 3.66 | Lys214 (O) | | | | | | |

**Salt Bridges**

| LaG41 | Dist. (Å) | GFP | LaG21 | Dist. (Å) | GFP | LaG19 | Dist. (Å) | GFP |
|---|---|---|---|---|---|---|---|---|
| Asp111 (OD2) | 3.55 | Lys45 (NZ) | Arg100 (NH1) | 3.31 | Glu32 (OE1) | Asp111 (OD1) | 3.94 | Lys45 (NZ) |
| Asp111 (OD1) | 2.16 | Lys45 (NZ) | Arg100 (NH2) | 3.99 | Glu32 (OE1) | Asp111 (OD2) | 2.56 | Lys45 (NZ) |
| Asp58 (OD1) | 3.60 | Arg215 (NH1) | Arg100 (NH1) | 3.18 | Glu32 (OE2) | Asp58 (OD1) | 3.69 | Arg215 (NH1) |
| Asp58 (OD2) | 2.57 | Arg215 (NH1) | Arg100 (NH2) | 2.42 | Glu32 (OE2) | Asp58 (OD2) | 2.68 | Arg215 (NH1) |
| Asp58 (OD1) | 3.35 | Arg215 (NH2) | Arg102 (NE) | 3.59 | Glu32 (OE2) | Asp58 (OD1) | 2.80 | Arg215 (NH2) |
| Asp58 (OD2) | 3.73 | Glu32 (OE1) | Arg102 (NH1) | 3.11 | Glu32 (OE2) | Asp58 (OD2) | 3.26 | Arg215 (NH2) |
| Arg100 (NH1) | 3.51 | Glu32 (OE1) | Asp111 (OD2) | 2.58 | Lys45 (NZ) | Arg100 (NH1) | 2.91 | Glu32 (OE1) |
| Arg100 (NH2) | 3.61 | Glu32 (OE1) | Asp58 (OD1) | 3.65 | Arg215 (NH1) | Arg100 (NH2) | 3.50 | Glu32 (OE1) |
| Arg102 (NE) | 2.55 | Glu32 (OE1) | Asp58 (OD2) | 2.56 | Arg215 (NH1) | Arg100 (NH1) | 3.65 | Glu32 (OE2) |
| Arg102 (NH1) | 4.00 | Glu32 (OE1) | Asp58 (OD1) | 2.85 | Arg215 (NH2) | Arg100 (NH2) | 2.66 | Glu32 (OE2) |
| Arg102 (NH2) | 3.47 | Glu32 (OE1) | Asp58 (OD2) | 3.24 | Arg215 (NH2) | | | |
| Arg100 (NH1) | 3.04 | Glu32 (OE2) | | | | | | |

**Supplementary Table 5.** Binding kinetics of Nanobody point variants relating to Figures 3 and 4 of main text.

| | Nanobody | $K_{on}$ ($M^{-1}$ $s^{-1}$) | $K_{off}$ ($s^{-1}$) | KD (M) | $Chi^2$ ($RU^2$) |
|---|---|---|---|---|---|
| **Paratope point variants** | $LaG43_{N78R}$ | 3.92E+06 | 1.22E-02 | 3.11E-09 | 3.71E-01 |
| | $LaG43_{K80R}$ | 1.82E+06 | 1.74E-02 | 9.00E-09 | 1.12 |
| | $LaG43_{N81R}$ | 7.94E+06 | 1.60E-02 | 2.00E-09 | 5.3 |
| | $LaG19_{S102R}$ | 1.20E+07 | 6.80E-03 | 5.80E-10 | 2 |
| | $LaG19_{Q103V}$ | 2.97E+06 | 1.83E-02 | 6.14E-09 | 1.42 |
| | $LaG19Q_{103I}$ | 1.62E+06 | 9.24E-02 | 5.70E-08 | 2.95E-01 |
| **FW point variants** | $LaG21_{L69F}$ | 2.50E+06 | 1.80E-03 | 7.30E-10 | 1.42 |
| | $LaG16_{E91K}$ | 2.02E+06 | 2.34E-03 | 1.15E-09 | 3.02E-01 |
| | $LaG24_{Q68K}$ | 1.87E+05 | 1.14E-03 | 6.09E-09 | 2.59E-01 |
| | $LaG24_{H85Q}$ | 3.68E+05 | 1.30E-03 | 3.52E-09 | 2.01E-01 |

**Supplementary Table 6.** Amino acid sequences of wildtype nanobody and GFP constructs used in this study

| Protein | Amino acid sequence |
|---|---|
| LaG16 | GSMAQVQLVESGGRLVQAGDSLRLSCAASGRTFSTSAMAWFRQAPGREREFVAAITW<br>TVGNTILGDSVKGRFTISRDRAKNTVDLQMDNLEPEDTAVYYCSARSRGYVLSVLRSVD<br>SYDYWGQGTQVTVSGLEVLFQGPSLEHHHHHH |
| LaG19 | GSMAQVQLVESGGGLVQAGGSLRLSCAASGPTGAMAWFRQAPGKEREFVGGISRSG<br>TDTYYVDSVKGRFTIDRDNAKNTVYLQMNSLKPEDTAVYYCAARRSQILFTSRTDYEFW<br>GQGTQVTVSGLEVLFQGPSLEHHHHHH |
| LaG21 | GSMAQVQLVESGGGLVQAGGSLRLSCAASGPTGAMAWFRQAPGMEREFVGGISGSE<br>TDTYYADFVKGRLTVDRDNVKNTVDLQMNSLKPEDTAVYYCAARRRVTLFTSRADYDF<br>WGQGTQVTVSGLEVLFQGPSLEHHHHHH |
| LaG24 | GSMADVQLVESGGGLVQPGGSLRLSCAASGEIASIIAIGWYRQAPGKQRESVALITRSG<br>MITYGDSAQGRFTISRDDAKNTVYLHMDDLVPEDTAVYYCNAKKVSFGDYWGQGTQV<br>TVSGLEVLFQGPSLEHHHHHH |
| LaG35 | GSMADVQLVESGGGLVQAGGSLRLSCTVSGRTFSNYAMGWFRQAPGKEREFVAGIS<br>WTGGHTLYTDSVKGRFTISRDNAKNTVYLQMNSLKPEDTALYYCAADRAADFFAQRDE<br>YDYWGQGTQVTVSGLEVLFQGPSLEHHHHHH |
| LaG41 | GSMADVQLVESGGGLVQAGGSLRLSCAASGPTGAMAWFRQAPGKEREFVGGISGSE<br>TDTYYVDSVKGRFTVDRDNVKNTVYLQMNSLKPEDTAVYYCAARRRITLFTSRTDYDF<br>WGRGTQVTVGLEVLFQGPSLEHHHHHH |
| LaG43 | GSMADVQLVESGGGLVQPGGSLRLSCEASGGAFSTVAMGWFRQAPGKEREFVGAIT<br>WTAGSTYYADSAKGRFTISRDNAKNTVHLQMNSLKPEDTAVYYCAQRVRGFFGPLRTT<br>PSWYEYWGQGTQVTVSGLEVLFQGPSLEHHHHHH |
| LaG16 | GSMAQVQLVESGGRLVQAGDSLRLSCAASGRTFSTSAMAWFRQAPGREREFVAAITW<br>TVGNTILGDSVKGRFTISRDRAKNTVDLQMDNLEPEDTAVYYCSARSRGYVLSVLRSVD<br>SYDYWGQGTQVTVSGLEVLFQGPSLEHHHHHH |
| green fluorescent protein (GFP) | MASKGEELFTGVVPILVELDGDVNGHKFSVSGEGEGDATYGKLTLKFICTTGKLPVPWP<br>TLVTTFGYGVQCFARYPDHMKQHDFFKSAMPEGYVQERTIFFKDDGNYKTRAEVKFEG<br>DTLVNRIELKGIDFKEDGNILGHKLEYNYNSHNVYIMADKQKNGIKVNFKIRHNIEDGSVH<br>LADHYQQNTPIGDGPVLLPDNHYLSTQSALSKDPNEKRDHMVLLEFVTAAGITHGMDEL<br>YKGLEVLFQGPSHHHHHH |

**Supplementary Table 7.** Amino acid sequences of nanobody variant constructs used in Figures 3 and 4 in main text and Supplementary Fig. 5.

| Nanobody | Amino acid sequence |
|---|---|
| LaG43$_{N78R}$ | GSMADVQLVESGGGLVQPGGSLRLSCEASGGAFSTVAMGWFRQAPGKEREFVGA<br>ITWTAGSTYYADSAKGRFTISRDR$_{78}$AKNTVHLQMNSLKPEDTAVYYCAQRVRGFFG<br>PLRTTPSWYEYWGQGTQVTVSGLEVLFQGPSLEHHHHHH |
| LaG43$_{K80R}$ | GSMADVQLVESGGGLVQPGGSLRLSCEASGGAFSTVAMGWFRQAPGKEREFVGA<br>ITWTAGSTYYADSAKGRFTISRDNAR$_{80}$NTVHLQMNSLKPEDTAVYYCAQRVRGFFG<br>PLRTTPSWYEYWGQGTQVTVSGLEVLFQGPSLEHHHHHH |
| LaG43$_{N81R}$ | GSMADVQLVESGGGLVQPGGSLRLSCEASGGAFSTVAMGWFRQAPGKEREFVGA<br>ITWTAGSTYYADSAKGRFTISRDNAKR$_{81}$TVHLQMNSLKPEDTAVYYCAQRVRGFFG<br>PLRTTPSWYEYWGQGTQVTVSGLEVLFQGPSLEHHHHHH |
| LaG19$_{S102R}$ | GSMAQVQLVESGGGLVQAGGSLRLSCAASGPTGAMAWFRQAPGKEREFVGGISR<br>SGTDTYYVDSVKGRFTIDRDNAKNTVYLQMNSLKPEDTAVYYCAARRR$_{102}$QILFTSR<br>TDYEFWGQGTQVTVSGLEVLFQGPSLEHHHHHH |
| LaG19$_{Q103V}$ | GSMAQVQLVESGGGLVQAGGSLRLSCAASGPTGAMAWFRQAPGKEREFVGGISR<br>SGTDTYYVDSVKGRFTIDRDNAKNTVYLQMNSLKPEDTAVYYCAARRSV$_{103}$ILFTSR<br>TDYEFWGQGTQVTVSGLEVLFQGPSLEHHHHHH |
| LaG19$_{Q103I}$ | GSMAQVQLVESGGGLVQAGGSLRLSCAASGPTGAMAWFRQAPGKEREFVGGISR<br>SGTDTYYVDSVKGRFTIDRDNAKNTVYLQMNSLKPEDTAVYYCAARRSI$_{103I}$LFTSRT<br>DYEFWGQGTQVTVSGLEVLFQGPSLEHHHHHH |
| LaG21$_{L69F}$ | GSMAQVQLVESGGGLVQAGGSLRLSCAASGPTGAMAWFRQAPGMEREFVGGISG<br>SETDTYYADFVKGRF$_{69}$TVDRDNVKNTVDLQMNSLKPEDTAVYYCAARRRVTLFTSR<br>ADYDFWGQGTQVTVSGLEVLFQGPSLEHHHHHH |
| LaG16$_{E91K}$ | GSMAQVQLVESGGRLVQAGDSLRLSCAASGRTFSTSAMAWFRQAPGREREFVAAI<br>TWTVGNTILGDSVKGRFTISRDRAKNTVDLQMDNLK$_{91}$PEDTAVYYCSARSRGYVLS<br>VLRSVDSYDYWGQGTQVTVSGLEVLFQGPSLEHHHHHH |
| LaG24$_{Q68K}$ | GSMADVQLVESGGGLVQPGGSLRLSCAASGEIASIIAIGWYRQAPGKQRESVALITR<br>SGMITYGDSAK$_{68}$GRFTISRDDAKNTVYLHMDDLVPEDTAVYYCNAKKVSFGDYWG<br>QGTQVTVSGLEVLFQGPSLEHHHHHH |
| LaG24$_{H85Q}$ | GSMADVQLVESGGGLVQPGGSLRLSCAASGEIASIIAIGWYRQAPGKQRESVALITR<br>SGMITYGDSAQGRFTISRDDAKNTVYLQ$_{85}$MDDLVPEDTAVYYCNAKKVSFGDYWG<br>QGTQVTVSGLEVLFQGPSLEHHHHHH |
| GFP$_{E17A}$ | MASKGEELFTGVVPILVA$_{17}$LDGDVNGHKFSVSGEGEGDATYGKLTLKFICTTGKLPV<br>PWPTLVTTFGYGVQCFARYPDHMKQHDFFKSAMPEGYVQERTIFFKDDGNYKTRA<br>EVKFEGDTLVNRIELKGIDFKEDGNILGHKLEYNYNSHNVYIMADKQKNGIKVNFKIR<br>HNIEDGSVHLADHYQQNTPIGDGPVLLPDNHYLSTQSALSKDPNEKRDHMVLLEFVT<br>AAGITHGMDELYKGLEVLFQGPSHHHHHH |
| GFP$_{E32A}$ | MASKGEELFTGVVPILVELDGDVNGHKFSVSGA$_{32}$GEGDATYGKLTLKFICTTGKLPV<br>PWPTLVTTFGYGVQCFARYPDHMKQHDFFKSAMPEGYVQERTIFFKDDGNYKTRA<br>EVKFEGDTLVNRIELKGIDFKEDGNILGHKLEYNYNSHNVYIMADKQKNGIKVNFKIR<br>HNIEDGSVHLADHYQQNTPIGDGPVLLPDNHYLSTQSALSKDPNEKRDHMVLLEFVT<br>AAGITHGMDELYKGLEVLFQGPSHHHHHH |

Residues mutated to create variants are marked in red.

**Supplementary Table 8.** Amino acid sequences of wildtype and variant anti-SARS-CoV-2 nanobody constructs used in Table 1.

| Nanobody | Amino acid sequence |
|---|---|
| S1-8 | MAQVQLVESGGGLVQAGGSLRLSCAASGRVLSSYVMGWFRQAPGKEREFVAAIRWNGGSTFYADSVKGRLTISRDNAKNTVYLQMNSLKPEDTAAYYCAASPRTMYDATYYYSRTSYDYWGQGTQVTVS |
| S1-8$_{L70F}$ | MAQVQLVESGGGLVQAGGSLRLSCAASGRVLSSYVMGWFRQAPGKEREFVAAIRWNGGSTFYADSVKGRF$_{70}$TISRDNAKNTVYLQMNSLKPEDTAAYYCAASPRTMYDATYYYSRTSYDYWGQGTQVTVS |
| S1-28 | MAQVQLVESGGGLVQAGGSLKLSCAVSGRTLSSYVMGWFRQAPGKEREFVAAIRWNGGSTFYADSVQGRLTISRDNAKNMVYLQMDSLKPEDTAAYYCAASPRTMYYASYYYTRTSYDYWGQGTQVTVS |
| S1-28$_{L70F}$ | MAQVQLVESGGGLVQAGGSLKLSCAVSGRTLSSYVMGWFRQAPGKEREFVAAIRWNGGSTFYADSVQGRF$_{70}$TISRDNAKNMVYLQMDSLKPEDTAAYYCAASPRTMYYASYYYTRTSYDYWGQGTQVTVS |
| S1-36 | MAQVQLVESGGGLVQAGGSLRVSCAASERILSSYVMGWFRQAPGKEREFVAAIRWNGGSTFYADSVKGRLTISRDNAKNTVYLQMNSLKPEDTAVYYCAASPRTMYLASYYYHRTSYDYWGQGTQVTVS |
| S1-36$_{L70F}$ | MAQVQLVESGGGLVQAGGSLRVSCAASERILSSYVMGWFRQAPGKEREFVAAIRWNGGSTFYADSVKGRF$_{70}$TISRDNAKNTVYLQMNSLKPEDTAVYYCAASPRTMYLASYYYHRTSYDYWGQGTQVTVS |
| S1-40 | MAQVQLVESGGGLVQVGGSLRLSCAASGRTFSVTGMGWFRQAPGKEREFVATINWRGDYTDYADSVKGRFTISRDNAKSTVYLQMNSLSSEDTAVYYCAGDRRGYGDSRSVVYDYWGQGTQVTVS |
| S1-40$_{S89K}$ | MAQVQLVESGGGLVQVGGSLRLSCAASGRTFSVTGMGWFRQAPGKEREFVATINWRGDYTDYADSVKGRFTISRDNAKSTVYLQMNSLK$_{89}$SEDTAVYYCAGDRRGYGDSRSVVYDYWGQGTQVTVS |
| S1-RBD-14 | MAQVQLVESGGGLVQAGGSVRLSCAASGRSFSINPMGWFRQAPGKEREFVAAISWSGSKTVYVDSVKGRFSISRDNAKNTVYLQMNSLEPEDTAVYHCAVASRGPVYGANYVPGPHEYEYWGQGTQVTVS |
| S1-RBD-14$_{E89K}$ | MAQVQLVESGGGLVQAGGSVRLSCAASGRSFSINPMGWFRQAPGKEREFVAAISWSGSKTVYVDSVKGRFSISRDNAKNTVYLQMNSLK$_{89}$PEDTAVYHCAVASRGPVYGANYVPGPHEYEYWGQGTQVTVS |
| S1-RBD-42 | MAQVQLVESGGGLVQAGGSLRLSCVVSGSTFSNTFMDWYRQAPGKQREFVATISSGGTTNYAVFVKGRFTISRDGAKNTVYLQMNSLEPEDTAVYYCHAALPIGGDYWGQGTQVTVS |
| S1-RBD-42$_{E88K}$ | MAQVQLVESGGGLVQAGGSLRLSCVVSGSTFSNTFMDWYRQAPGKQREFVATISSGGTTNYAVFVKGRFTISRDGAKNTVYLQMNSLK$_{88}$PEDTAVYYCHAALPIGGDYWGQGTQVTVS |
| S1-14 | MAQVQLVESGGGLVQAGGSLRLSCAASGSTFSRLSMGWYRQVPGKQRELVARILPLGGPYYRDFVQGRFTISRDNVKNMLYLQMNSLKPEDTAVYYCNRAPFGTAWDGPDNYDYWGQGTQVTVS |
| S1-14$_{Q66K}$ | MAQVQLVESGGGLVQAGGSLRLSCAASGSTFSRLSMGWYRQVPGKQRELVARILPLGGPYYRDFVK$_{66}$GRFTISRDNVKNMLYLQMNSLKPEDTAVYYCNRAPFGTAWDGPDNYDYWGQGTQVTVS |
| S1-RBD-31 | MGQVQLVESGGGLVQAGDSLRLSCTASGRSFSTNAMGWFRQAPGKEREFVSAISWSSGTTYYSDSVQGRFTISGDNAKNTVYLQMNELKPDDTAVYYCTLRSQFNAYAWTTKYAYDYWGQGTQVTVS |
| S1-RBD-31$_{Q67K}$ | MGQVQLVESGGGLVQAGDSLRLSCTASGRSFSTNAMGWFRQAPGKEREFVSAISWSSGTTYYSDSVK$_{67}$GRFTISGDNAKNTVYLQMNELKPDDTAVYYCTLRSQFNAYAWTTKYAYDYWGQGTQVTVS |

Residues mutated to create framework variants are marked in red.
